# Testing concordance and conflict in spatial replication of landscape genetics inferences

**DOI:** 10.1101/2021.06.21.449301

**Authors:** Van Wishingrad, Robert C. Thomson

## Abstract

The field of landscape genetics relates habitat features and genetic information to infer dispersal and genetic connectivity between populations or individuals distributed across a landscape. Such studies usually focus on a small portion of a species range, and the degree to which these geographically restricted results can be extrapolated to different areas of a species range remains poorly understood. Studies that have focused on spatial replication in landscape genetics processes either evaluate a small number of sites, are informed by a small set of genetic markers, analyze only a small subset of environmental variables, or implement models that do not fully explore parameter space. Here, we used a broadly distributed ectothermic lizard (*Sceloporus occidentalis,* Western Fence lizard) as a model species to evaluate the full role of topography, climate, vegetation, and roads on dispersal and genetic differentiation. We conducted landscape genetics analyses in five areas within the Sierra Nevada mountain range, using thousands of ddRAD genetic markers distributed across the genome, implemented in the landscape genetics program ResistanceGA. Across study areas, we found a great deal of consistency in the variables impacting genetic connectivity, but also noted site-specific differences in the factors in each study area. High-elevation colder areas were consistently found to be barriers to gene flow, as were areas of high ruggedness and slope. High temperature seasonality and high precipitation during the winter wet season also presented a substantial barrier to gene flow in a majority of study areas. The effect of other landscape variables on genetic differentiation was more idiosyncratic and depended on specific attributes at each site. Vegetation type was found to substantially affect gene flow only in the southernmost Sequoia site, likely due to a higher proportion of desert habitat here, thereby fragmenting habitats that have lower costs to dispersal. The effect of roads also varied between sites and may be related to differences in road usage and amount of traffic in each area. Across study areas, canyons were always substantially implicated as facilitators to dispersal and key features linking populations and maintaining genetic connectivity across landscapes. We emphasize that spatial data layers are complex and multidimensional, and a careful consideration of associations between variables is vital to form sound conclusions about the critical factors affecting dispersal and genetic connectivity across space.

## INTRODUCTION

Landscape genetics theory predicts that the degree of genetic differentiation experienced by populations distributed across a landscape is due to habitat, climate, and topographic factors affecting rates of allele migration across these features (Manel *et al.* 2003; Storfer *et al.* 2010; Manel & Holderegger 2013; Balkenhol *et al.* 2016). Implicit in this theory is the assumption that these barriers to gene flow can be accurately assessed and used to predict genetic connectivity in novel areas where similar landscape variables are found. Until very recently, nearly all landscape genetics studies related gene flow estimates to landscape structure in a single species within a single landscape (Segelbacher *et al.* 2010). As the field has evolved, however, researchers have realized the importance of understanding the degree to which landscape genetics studies can be extrapolated, or replicated, across different landscapes (Short Bull *et al.* 2011; Richardson 2012; Trumbo *et al.* 2013; Zancolli *et al.* 2014; Selkoe *et al.* 2016; Villemey *et al.* 2016; Burgess & Garrick 2020). For example, Short Bull *et al.* (2011) conducted a landscape genetics study on black bears (*Ursus americanus*) in 12 study areas that varied in extent size, landscape composition, variability, and complexity. They found that barriers to gene flow varied among regions, and gene flow was related to the amount of variation in landscape elements across the landscape. In another study, Trumbo *et al.* (2013) investigated differences in landscape composition in several parts of the range of the Cope’s giant salamander (*Dicamptodon copei*) and found that stream and river networks in the northern portion of the range facilitated gene flow, whereas the southeastern edge of the species’ range, facing limitations of heat and forest fragmentation, restricted gene flow. Similarly, Burgess & Garrick (2020) compared landscape genetics results between two areas for five landcover variables and found that in both study areas, agriculture was associated with resistance to gene flow, while wetlands had opposite effects on gene flow—facilitating gene flow in one area, while inhibiting gene flow in another. Finally, a recent review of seascape genomics studies revealed that when more variables are included in an analysis a smaller proportion of those variables are significant predictors of genetic connectivity, suggesting that a few drivers tend to dominate in marine systems (Selkoe *et al.* 2016). These studies have all contributed to our understanding of spatial concordance in landscape genetics. However, comparative studies of genomic-scale variation which harness methods that fully explore parameter space promise to provide a higher level of accuracy than previous approaches. Furthermore, evaluating all relevant environmental variables across a larger number of sites will help more fully explore the degree of concordance and conflict in landscape genetics processes across space.

The Sierra Nevada is an ideal setting to make comparisons between different areas that bear many resemblances in their environment. These include similar plant communities along the extent of the range, such as vegetation stratification along elevation gradients with similar communities in each strata (Schoenherr 1992). Along the western slopes, the lowest elevation characterized by foothill woodland and chaparral spanning from 150 m – 1000 m, followed by lower montane forest from 1000 m – 1800 m, mid-montane forest from 1000 m – 2000 m, upper montane forest from 2000 m – 2700 m, subalpine forest from 2700 m – to 3400 m, and the alpine zone generally above 3200 m (Schoenherr 1992). The eastern biotic zones are similar but occur at a higher elevation, presumably due to reduced precipitation on the eastern slope of the Sierra Nevada, and includes lower montane forest from 2000 m – 2700 m, upper montane forest from 2700 m – 3200 m, subalpine forest from 3200 m – 3500 m, and the alpine zone above 3500 m (Schoenherr 1992). The Sierra Nevada range also exhibits many similarities in terms of topographic variability and complexity along its extent. The geologic history of the Sierra Nevada is complex, but is the entirety of the Sierra Nevada mountain range is generally believed to have been formed by common geological processes (McPhee 1993), and bears many similarities in the structure of river canyons that span predominantly east-west across the latitudinal extent of the Sierra Nevada. Climate patterns are also largely similar across the Sierra Nevada range, characterized by cold, wet winters and warm, dry summers that often give way to afternoon thunderstorms mid- to late-summer (Schoenherr 1992). Precipitation tends to be highest (500 – 2000 mm/annually) as moisture enters the western slope from 1500 m – 2400 m, leading to a “rain shadow” effect with lower levels of precipitation (typically less than 600 mm/annually) in the eastern Sierra Nevada slopes (Schoenherr 1992). These similarities in environment make it possible to compare whether a species shows the same response to a common environment in different areas, helping to understand if genetic connectivity is a feature of the landscape (i.e., different barriers and facilitators to gene flow between areas), or of the species (i.e., similar barriers and facilitators to gene flow between areas).

Studies investigating landscape genetics processes would benefit from model species whose ecology is well understood and is widely distributed over an expanse of different habitat types, as these results may be more generalizable to other taxonomic groups inhabiting a range of environments. The Western Fence lizard (*Sceloporus occidentalis*) is a generalist lizard found across a diverse range of habitat types and elevation gradients from sea level to elevations of up to 3300 m spanning a wide-range of environmental gradients and widely distributed across Western North America (Stebbins 2003). Western Fence lizards are also found in a wide variety of habitats, including grasslands, chaparral, sagebrush, woodland, and coniferous forests. They are generally excluded from harsh deserts and elevations above 3300 m, making these habitats potentially strong barriers to gene flow. We predict that high-elevation, colder, regions will restrict gene flow, along with steep and rugged canyons, harsh desert regions, water bodies, and roads, and results will be largely similar across study areas.

Here, we conducted five large-scale landscape genetics studies, and compared the inferred dispersal costs through areas that differ in elevation, temperature, precipitation, topographic features, vegetation types, and roads. Each of the study areas are, to the extent that was possible, composed of similar habitats, elevation gradients, and are of identical size (50 km × 50 km extents) (Figure 1). This comparison will inform whether the same landscape elements are impediments or facilitators to gene flow (e.g., woodlands, temperature and/or moisture gradients, topographic relief) in distinct study areas and should shed light on the spatial generalizability of landscape genetics inferences in Western Fence lizards. Of particular interest is the degree of concordance or conflict in the relative dispersal costs through different habitat types in each of the study areas. If landscape effects on gene flow are conserved within the species, then the same general results should be found in different regions. If not, then it suggests these landscape genetics inferences may be population-specific due to unique adaptations that change the way they experience the landscape (e.g., local adaptation), or landscapes themselves are fundamentally different at different latitudes (e.g., exposed areas at high latitude are physiologically tolerable, while at low latitude they act as barriers to migration), making it difficult to extrapolate landscape genetics inferences across space (Schoville et al. 2011).

**Figure 1.**
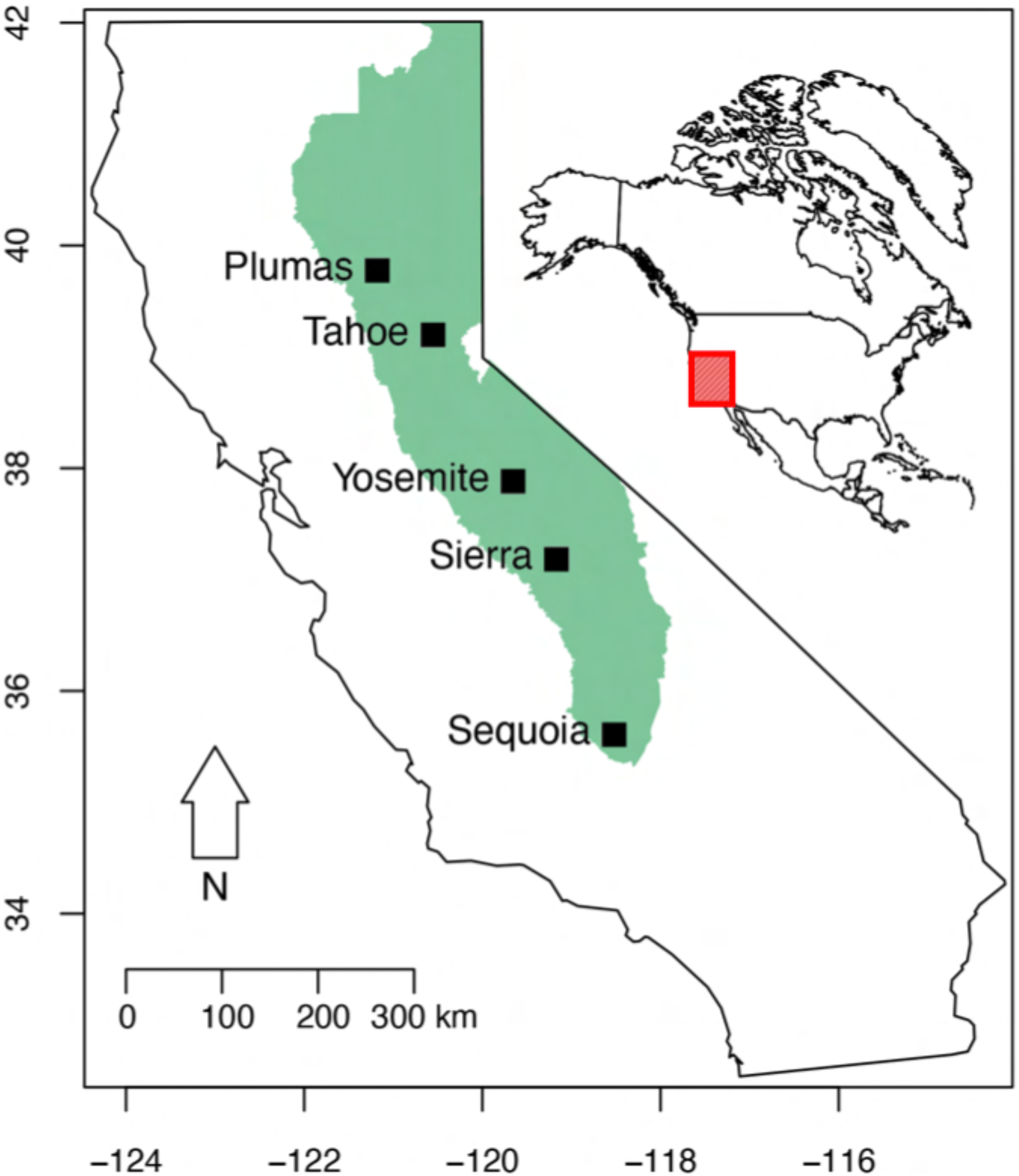
Overview map of the five study areas situated within the Sierra Nevada Conservancy (green shaded area) and their location in North American continent (red shaded area).

## METHODS

### LIZARD COLLECTION

We captured lizards, either by hand or by lizard nooses, and recorded Global Positioning System (GPS) coordinates with a Garmin^®^ GPS 60 at the site of capture. We sampled lizards from June through August in 2016 and 2018 in California, USA over a latitudinal range of >900 km and an elevation range of 100 – 2800 m (Figure 1). Our focus was on the Sierra Nevada mountain range, which encompasses a large portion of environmental heterogeneity across this species’ range. For approximately half of the individuals collected, we followed a standard euthanasia protocol based on Conroy et al. (2009), with modifications specific to *Sceloporus*, and prepared specimens as vouchers, preserving liver samples in 95% ethanol for DNA extraction. All voucher specimens and genetic samples are now deposited at the Natural History Museum of Los Angeles County, section of Herpetology (see Supplemental Information for details including voucher numbers). For the other half of lizards collected, we removed a <5 mm portion of the tail’s distal portion and preserved the tissue in 95% ethanol for DNA extraction. We also obtained tissue samples from museum specimens at the Museum of Vertebrate Zoology at University of California, Berkeley; the California Academy of Sciences; the Yale Peabody Museum of Natural History; and from ongoing collaborations.

### ddRAD LIBRARY PREPARATION AND GENOMIC SEQUENCING

We followed Peterson *et al.*’s (2012) method for genomic library preparation, with some modifications. For each individual, we extracted high-molecular weight genomic DNA using a standard phenol-chloroform extraction protocol (Tsai *et al.* 2019). We measured DNA concentrations using a Qubit fluorometer, and for each sample, we digested 0.5 μg of DNA for 3 hours with SbfI and NIaIII restriction enzymes. We then purified these fragments with Agencourt AMPure beads before ligation of barcoded Illumina adaptors onto the fragmented ends. All barcodes differed by at least two base pairs to reduce duplexing error rates. We then pooled equimolar amounts of each sample before conducting size selection using a BluePippin Prep to select fragments between 400 and 550 bp in length. We used proofreading Taq and Illumina’s index primers for final library amplification for 8-10 cycles to reduce PCR bias. We quantified the final library concentration using a Qubit fluorometer at high sensitivity. We packed samples on dry ice then sent them to the Vincent J. Coates Genomics Sequencing Lab at UC Berkeley for sequencing. Quantitative PCR was used to determine concentration of adapter-associated fragments, and a BioAnalyzer run was used to confirm fragment sizes as a quality-control measure prior to sequencing. Final libraries were sequenced (100-bp or 150-bp, single-end runs) in Illumina HiSeq 4000 or NovaSeq SP lanes for a total of four sequencing lanes.

### BIOINFORMATICS PROTOCOLS FOR SNP DATA

We inspected raw Illumina reads for sample quality using FastQC (Andrews 2010). We demultiplexed our data using STACKS v1.48 (Catchen *et al.* 2013), and included steps to rescue barcodes with at most one mismatch, clean data, and remove any read with an uncalled base, and truncate all reads to 95 bases (due to variable-length sequences from using both 100-bp and 150-bp sequencing platforms). Prior to analysis, we concatenated sequencing data for the same individual sequenced on different lanes. We used a reference-based approach in both the STACKS (Catchen *et al.* 2013) population-based and ipyrad (Eaton & Overcast 2020) individual-based analyses. Compared to de-novo assembly approaches, reference-based approaches have much lower error rates, higher accuracy, and less bias than do de-novo approaches (Rochette & Catchen 2017). We used the annotated *S. occidentalis* genome published by Harris *et al.* (2015) as the reference genome for the analysis.

### IPYRAD ANALYSIS PROTOCOL

We followed the analysis recommendations and default settings specified by the program authors with some modifications specific to our datasets. We filtered and edited demultiplexed reads by removing reads with 5 or more low-quality base calls (Q<20), and trimmed bases from the 3’ end of reads if their quality scores fell below 20, which is 99% probability of a correct base call. Reads were then mapped to the reference genome using BWA (version 0.7.17-r1188) and clusters were aligned using MUSCLE (version 3.8.31) requiring a minimum depth of 6, which is the minimum depth at which a heterozygous base call can be distinguished from a sequencing error (ipyrad.readthedocs.io/en/0.9.79/). We jointly estimated heterozygosity and error rate by specifying a maximum of two alleles per site in each consensus sequence and removed alignments with a high proportion (5%) of heterozygous base calls, as poor alignments tend to have an excess of heterozygous sites. To remove poor alignments in repetitive regions in the final data set, we allowed for a maximum of 20% SNPs per locus, as well as removed alignments with more than 8 indels per locus. We set the maximum proportion of shared polymorphic sites in a locus to 0.5, as shared heterozygous sites across samples likely results from clustering of paralogs with fixed, rather than heterozygous, sites, and excluded any samples with fewer than 100,000 reads. We retained all loci shared by at least 75% of samples in the dataset for analysis. We retained 11,215 loci in the assembly with 25.9% missing sites in the final SNP matrix. We output an SNP-based formatted file of one randomly selected variable site per locus to calculate individual-based pairwise genetic distances calculated as a proportion of shared alleles for individuals collected over the extent of our sampling range (Figure 1). Sample sizes differ in each analysis and are detailed in the section specific for each study area.

### SPATIAL DATA LAYERS

We obtained land cover and vegetation data from the GAP/LANDFIRE National Terrestrial Ecosystems data sets (usgs.gov). These datasets are comprised of detailed vegetation and land cover data for 584 unique classes at several levels of classification and a 30-meter resolution scale. For the purpose of our analyses, we used the “Class” level classification, which partitions each unique class into 11 groups representing major land cover and vegetation categories. We obtained broad-scale climate data from the PRISM climate group (prism.oregonstate.edu). These data included: mean precipitation, mean temperature, and average monthly minimum and maximum temperatures over the most recent three full decades spanning the period 1981-2010 and are represented at a scale of 800-meter resolution. We also obtained bioclimatic data from the WorldClim database (worldclim.org) for variables hypothesized to be potentially important in limiting distribution and migration in reptiles. These data are represented at 30 seconds resolution (0.93 × 0.93 = 0.86 km2 at the equator) and include several aspects of temperature variation: maximum and minimum temperature during the warmest and coldest quarters (3-month period) and warmest and coldest months, along with two aspects of temperature variation, including temperature seasonality (standard deviation of temperature × 100) and temperature annual range (maximum temperature of warmest month – minimum temperature of coldest month). We also considered several aspects of precipitation variation in the environment, including maximum and minimum precipitation during the wettest and driest quarters (3-month period) and wettest and driest months, as well as precipitation seasonality (coefficient of variation in precipitation). We obtained elevation data captured by the Advanced Land Observing Satellite (ALOS) DAICHI-2 (eorc.jaxa.jp/ALOS/en/aw3d30/), which is the most precise global-scale elevation data we could obtain, at a resolution of 30-meters. Using the elevation data, we generated slope and ruggedness spatial layers in QGIS version 2.18 (qgis.org). Slope measures the angle of inclination of the terrain between adjacent cells, while ruggedness quantifies terrain heterogeneity as the change in elevation within a 3×3 cell grid (as described in (Riley *et al.* 1999). Finally, to investigate the effect of roads on genetic connectivity, we incorporated a spatial data layer from the United States Data Catalog of primary and secondary roads which includes U.S. Highways, State Highways, and/or County Highways (catalog.data.gov). Altogether, we retained 20 layers for the analyses: 1 vegetation layer, 4 PRISM climate layers, 11 WorldClim bioclimatic layers, 3 topographic layers, and 1 roads layer.

### SPATIAL DATA PROCESSING

We generated spatial layers representing the axes of greatest variation in the environment from all spatial layers comprised of continuous data, which included all temperature, precipitation, and topographic data layers. Specifically, we used R (version 3.6.1) and the package RStoolbox (version 0.2.6) to perform a PCA on stacks of raster layers. These principal component spatial layers represent different aspects of the environment in each spatial area as detailed in the results and supplemental materials. Because the relationships between variables differed between each of the study areas, we did not use a common set of PC axes to compare between regions, as a PC axis in one area may describe an environmental gradient that simply does not exist in another area. For each area, we retained the first three principal components—which was sufficient to describe >90% of the variation in the environment in each area—along with vegetation and roads layers, in each landscape genetics analysis. The vegetation category layer was converted to integer values 1-11, with each integer representing a distinct vegetation category. The roads layer was converted to a binary categorical layer specifying whether or not a road is found in the cell, where 0 indicates “no road” and 1 indicates “road”. In all cases, layers were converted to a common resolution of 0.0025 and 50 km-sq extents for a total size of 2,500 sq-km, each with 25,159 cells, and a cell size of ~1 km-sq.

### DESCRIPTION OF STUDY AREAS

#### SEQUOIA

Our Sequoia study site is situated at the southern end of the Sierra Nevada range (southwest corner of extent: −118.7500, 35.4211) in the Sequoia National Forest region (Figure 2) and includes 19 sampling sites (Figure 2a). This area includes portions of the Kern River Canyon, which is the largest drainage in the Sierra Nevada, and the entirety of Isabella Lake in the northeast. The study area is roughly bound by Breckenridge Mountain in the southwest, Piute Peak in the southeast, Glennville in the northwest, and Kernville in the northeast (Figure 2b). The landscape is predominantly desert and semi-desert composed of sagebrush scrubland and shrub-steppe especially in the lower-elevation areas surrounding Lake Isabella. Pinyon (*Pinus* spp.) and juniper (*Juniperus* spp.) woodlands are largely dominant in the southern parts of this extent, and present throughout the area at lower frequencies, increasing in density at higher elevations. Developed and other human-modified landscapes are present in some areas around Lake Isabella, and along the roads such as Highway 178 in the southwest, and Highway 155 in the northwest (Figure 2a-b). Agricultural and developed vegetation, recently disturbed and modified areas, and scrub, grasslands and barrens are present in smaller proportions in the landscape (Figure 2b).

**Figure 2.**
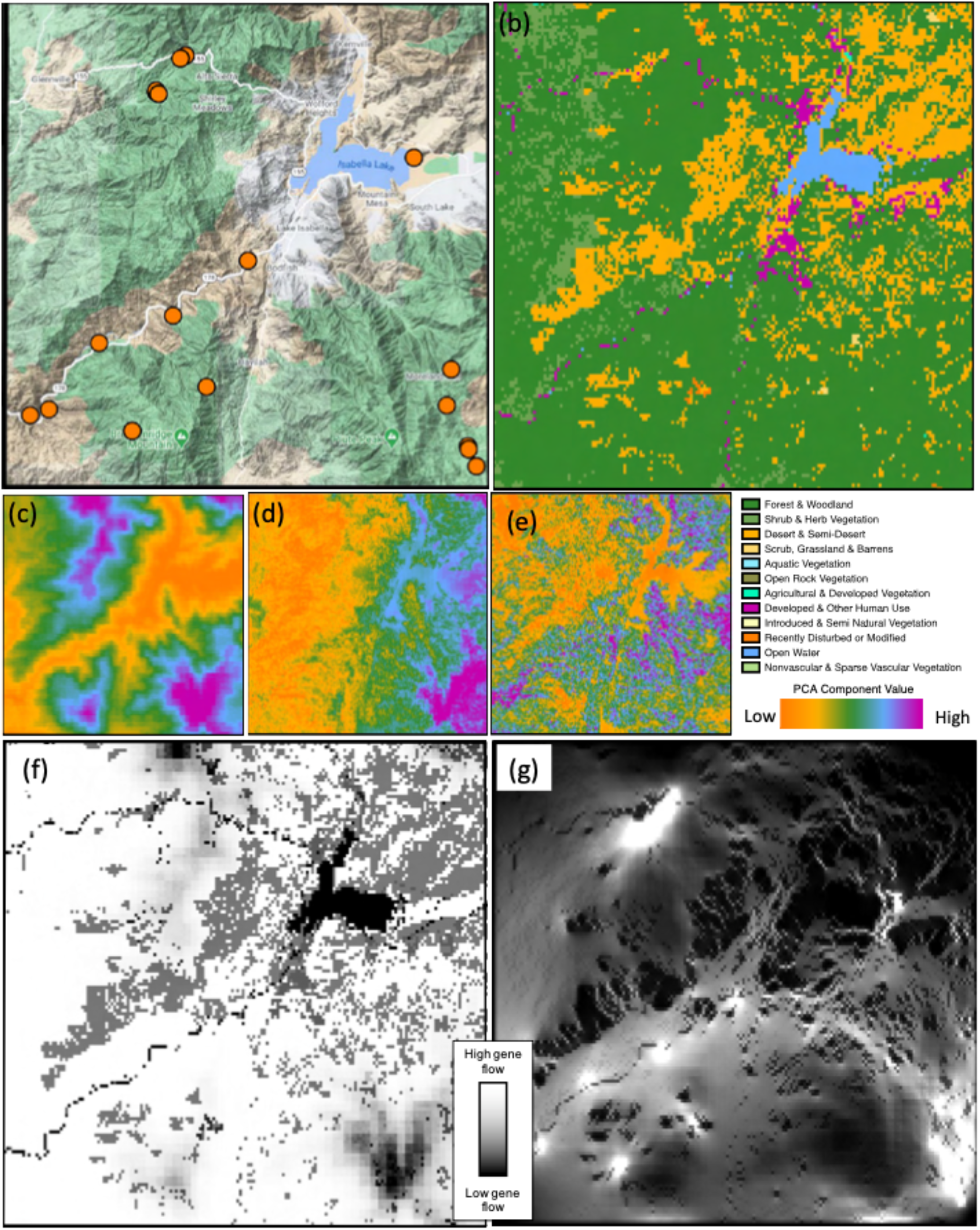
(a) Terrain map of the Sequoia region, where orange points indicate sampling locations (*n* = 19). Terrain map data © 2021 Google. (b) Vegetation map of the region, with each color representing a distinct vegetation category. (c) Heatmap of PCA component 1, where high values are associated with higher elevation regions with more precipitation and cooler temperatures, while low values are associated with lower elevation areas of higher temperatures. (d) Heatmap for PCA component 2, where high values are associated with areas that are more seasonal and experience more dry-season precipitation. (e) Heatmap for PCA component 3, where high values are associated with rugged areas of higher slope. (f) Optimized cost surface in the Sequoia region for the Comp1.Comp2.roads.vegetation surface. Light shading indicates low resistance values, while dark shading indicates areas of high resistance to gene flow. (g) Circuitscape output for the optimized cost surface in the Sequoia region with sampling sites as the focal nodes and the output raster Comp1.Comp2.roads.vegetation as the resistance surface. Light shading indicates high gene flow current, while dark shading indicates areas of low gene flow.

#### SIERRA

Our Sierra study site is located in the Sierra National Forest (southwest corner of extent: - 119.4001, 36.9887) in the southwest extent of the Sierra Nevada range (Figure 3) and includes 14 sampling sites (Figure 3a). This area includes Shaver Lake and Huntington Lakes near the center; Courtright Reservoir and Wishon Reservoir in the southeast; Florence Lake in the northeast; and Mammoth Pool Reservoir in the northwest (Figure 3a). The major read in this landscape is Highway 168, which runs from the southwest out of Fresno, California, through the town of Shaver Lake, and terminates in the town of Lakeshore on the northern shore of Huntington Lake. The landscape is predominantly forests and woodlands—including Jeffery pine (*Pinus jeffreyi*) and Ponderosa pine (*Pinus ponderosa*) woodlands as well as oak and other broadleaf woodlands, which includes several oak (*Quercus* spp.) species. Shrub and herb vegetation; desert and semi-desert; and scrub, grassland, and barren areas are interspersed throughout. Developed areas, introduced and semi-natural vegetation, and recently disturbed areas are present in lower frequencies (Figure 3b).

**Figure 3.**
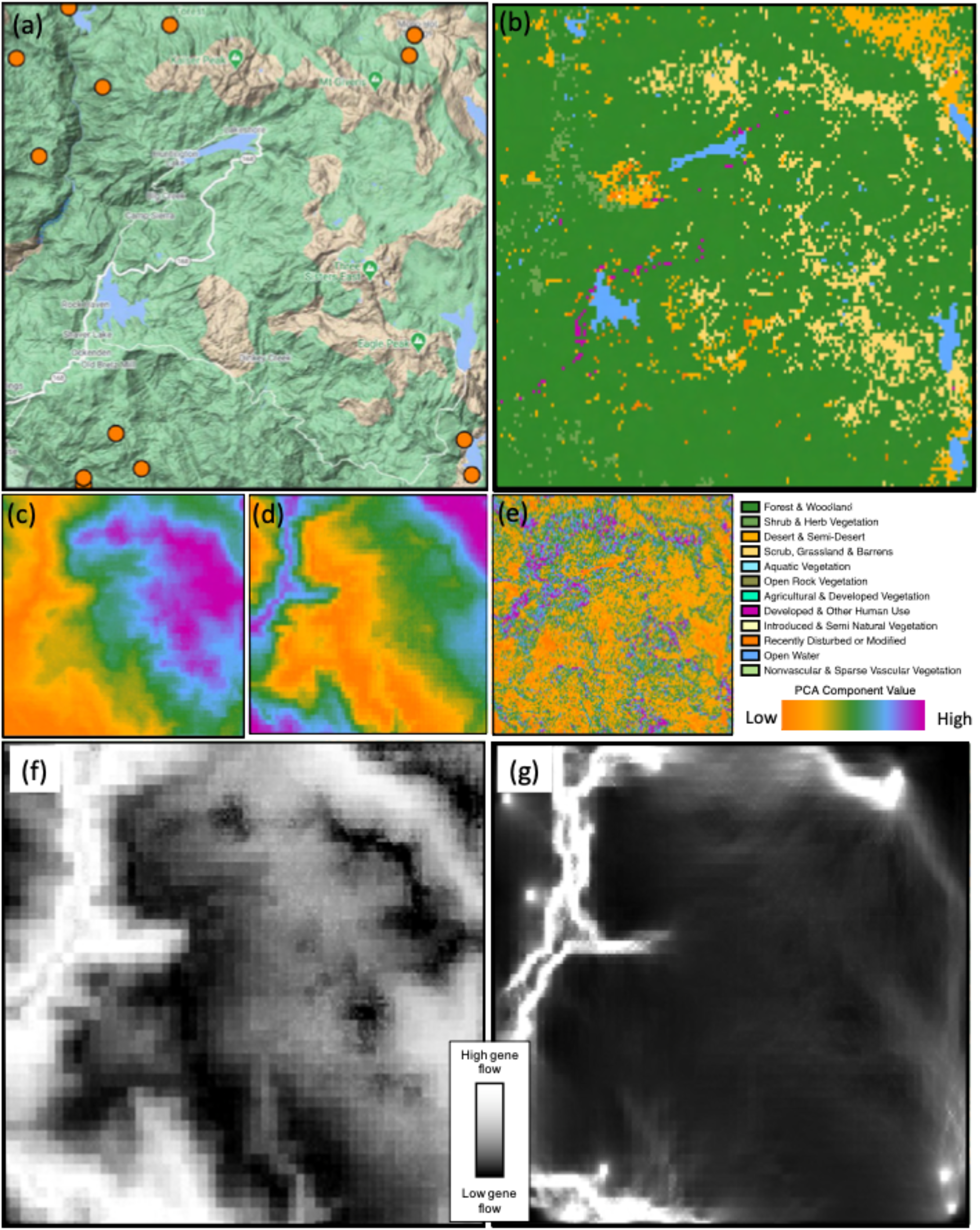
(a) Terrain map of the Sierra region, where orange points indicate sampling locations (*n* = 14). Terrain map data © 2021 Google. (b) Vegetation map of the region, with each color representing a distinct vegetation category. (c) Heatmap of PCA component 1, where high values are associated with higher elevation regions with more dry season precipitation and cooler temperatures, while low values are associated with lower elevation areas of higher temperatures. (d) Heatmap for PCA component 2, where high values are associated with areas of higher temperature seasonality and higher annual temperature range, while low values are associated with areas of high precipitation, especially wet season precipitation. (e) Heatmap for PCA component 3, where high values are associated with areas of high slope and ruggedness, while low values are associated with areas of low slope and ruggedness. (f) Optimized cost surface in the Sierra region for the Comp1.Comp2.Comp3 surface. Light shading indicates low resistance values, while dark shading indicates areas of high resistance to gene flow. (g) Circuitscape output for the optimized cost surface in the Sierra region with sampling sites as the focal nodes and the output raster Comp1.Comp2.Comp3 as the resistance surface. Light shading indicates high gene flow current, while dark shading indicates areas of low gene flow.

#### YOSEMITE

Our Yosemite study site is located in Yosemite National Park (southwest corner of extent: - 119.8851, 37.6988) and includes Yosemite Valley and Merced River on the southern extent, and Lake Eleanor and Hetch Hetchy Reservoir in the northwest (Figure 4) and includes 35 sampling sites (Figure 4a). Clouds Rest and Half Dome are located in the southeastern corner of the extent, Crane Flat in the southwest, and Piute Mountain and Slide Mountain in the northeast. Highway 140 is a major highway that runs along Merced River and through Yosemite Valley, and Highway 120 (Tioga Road) roughly bisects the lower third of the extent (Figure 4a). Yosemite National Park is a popular tourist destination, and both these roads are heavily used, especially in the summer months, averaging more than 4-5 million visitors annually in the past decade (irma.nps.gov). The landscape is dominated by forest and woodland habitat—including Jeffery pine, ponderosa pine woodlands, and mixed conifer woodlands; scrub, grassland, and barren areas that include sagebrush (*Artemesia* spp.) shrublands, dwarf shrublands, and sparsely vegetated tundra areas; and desert and semi-desert areas to a lesser degree. Developed and human-use areas are found near roads, and substantially in Yosemite Valley. Glacier-fed lakes and streams along with granite outcrops are found throughout the landscape (Figure 4b).

**Figure 4.**
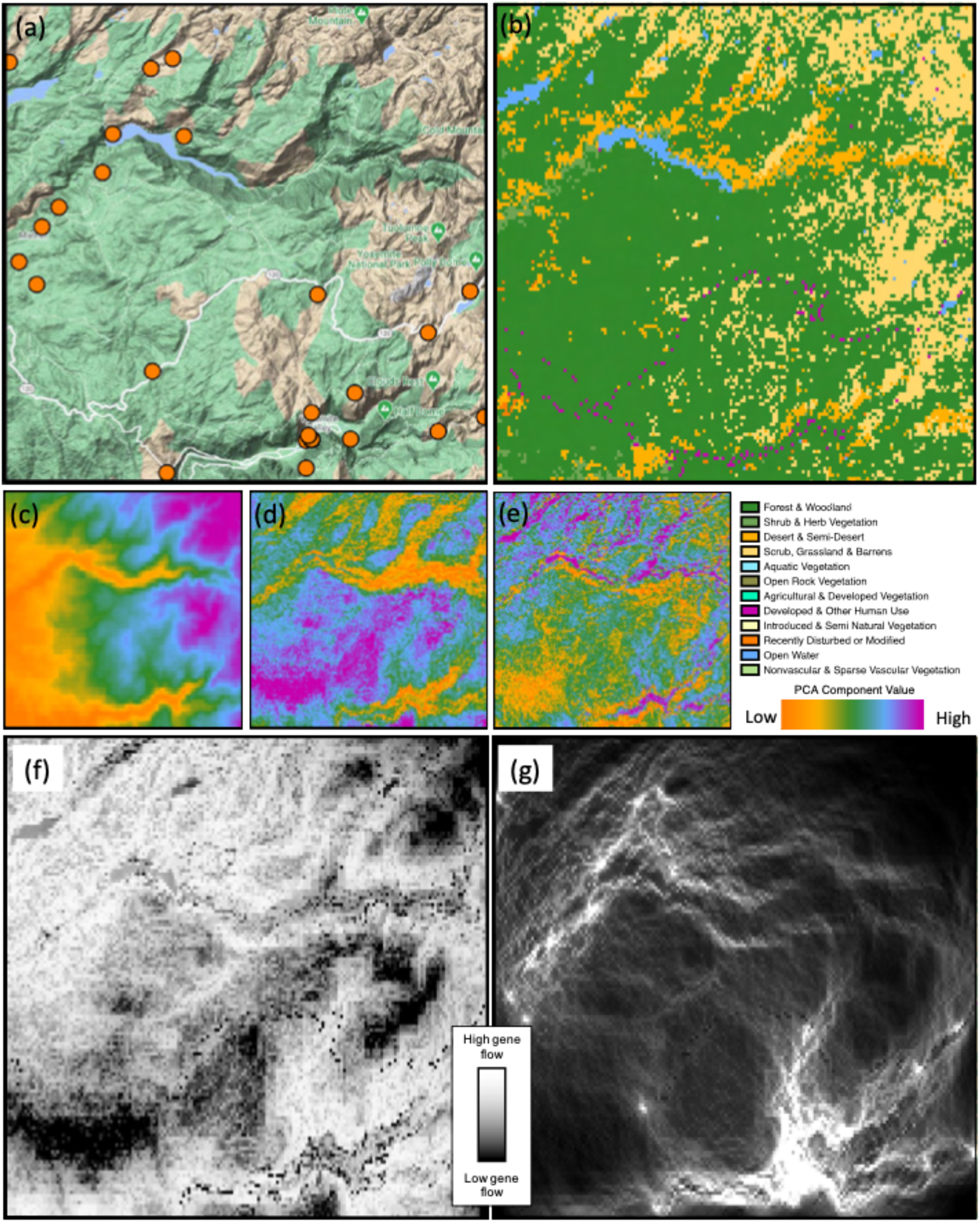
(a) Terrain map of the Yosemite region, where orange points indicate sampling locations (*n* = 35). Terrain map data © 2021 Google. (b) Vegetation map of the region, with each color representing a distinct vegetation category. (c) Heatmap of PCA component 1, where high values are associated with higher elevation regions with more dry season precipitation and cooler temperatures, while low values are associated with lower elevation areas of higher temperatures. (d) Heatmap for PCA component 2, where low values are associated with areas of higher topographic slope and ruggedness that also exhibit higher temperature seasonality and temperature annual range. (e) Heatmap for PCA component 3, where high values are associated with areas of high temperature seasonality and temperature annual range, while low values are associated with rugged areas of high slope. (f) Optimized cost surface in the Yosemite region for the Comp1.Comp2.Comp3.roads surface. Light shading indicates low resistance values, while dark shading indicates areas of high resistance to gene flow. (g) Circuitscape output for the optimized cost surface in the Yosemite region with sampling sites as the focal nodes and the output raster Comp1.Comp2.Comp3.roads layer as the resistance surface. Light shading indicates high gene flow current, while dark shading indicates areas of low gene flow.

#### TAHOE

Our Tahoe study site is located in Tahoe National Forest (southwest corner of extent: - 120.7876, 39.0138), roughly defined by Michigan Bluff in the southwestern corner; McKinstry Peak and Hell Hole Reservoir in the southeast; Highway 20 and Emerald Pools in the northwest; and Donner, Boreal Ridge, and the Sugar Bowl Resort in the northeast (Figure 5) and includes 39 sampling sites (Figure 5a). A major highway (Highway 80) runs across the northern extent of the range from east to west (Figure 5a). The landscape is dominated by forest and woodland habitat that includes Jeffery pine and ponderosa pine woodlands and mixed confer woodlands; smaller areas desert and semi desert featuring alpine dwarf-shrublands, forb meadows, and sparse grasslands; developed and human use areas along roads and recreational areas; and recently disturbed or modified landscapes. Other vegetation types are also found, though in lower frequencies (Figure 5b).

**Figure 5.**
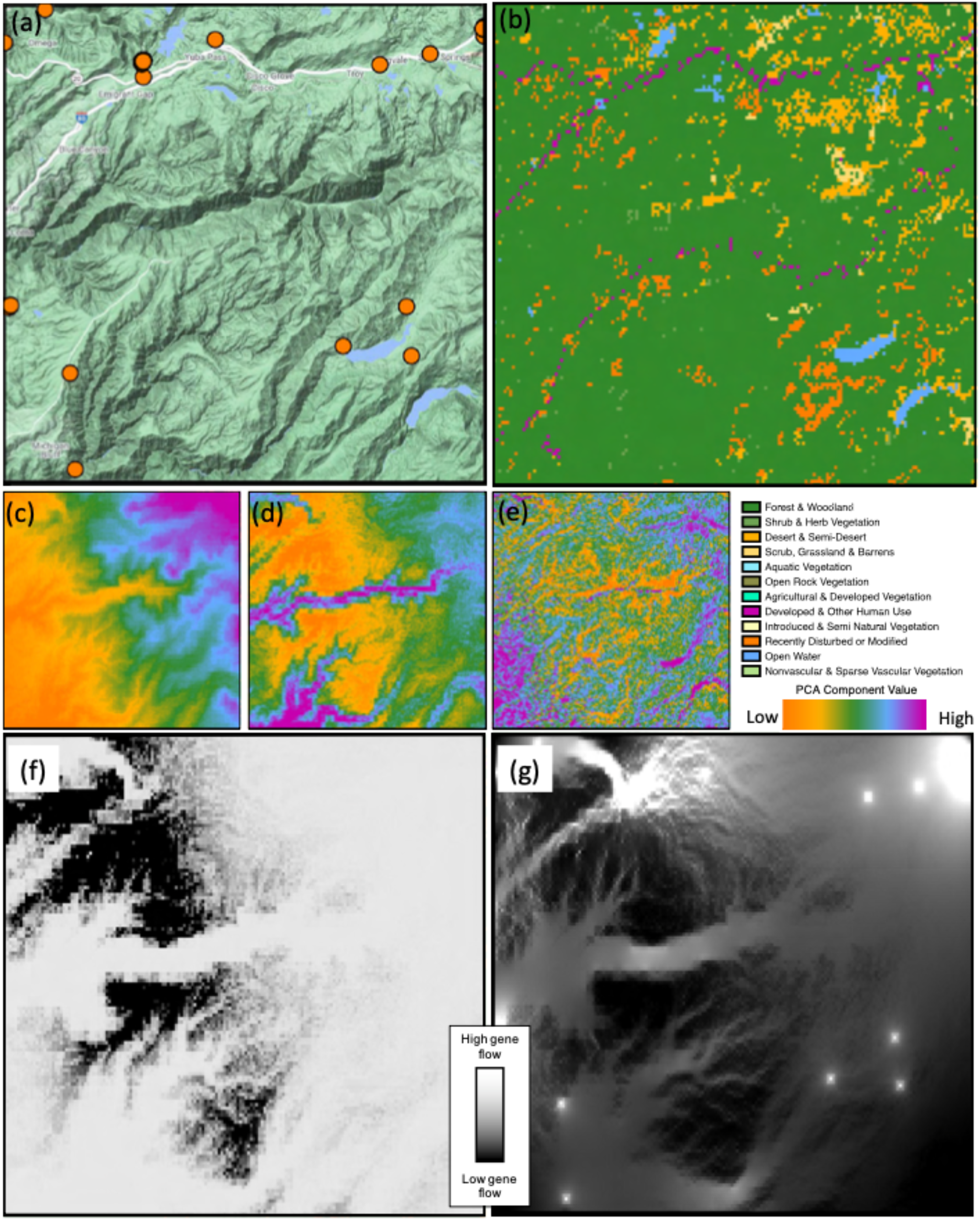
(a) Terrain map of the Tahoe region, where orange points indicate sampling locations (*n* = 39). Terrain map data © 2021 Google. (b) Vegetation map of the region, with each color representing a distinct vegetation category. (c) Heatmap of PCA component 1, where high values are associated with higher elevation regions with more dry season precipitation and cooler temperatures, while low values are associated with lower elevation areas of higher temperatures. (d) Heatmap for PCA component 2, where high values are associated with areas of high temperature seasonality, high temperature annual range, and rugged areas of high slope, while low values are associated with areas that receive more precipitation—especially wet season precipitation. (e) Heatmap for PCA component 3, where high values are those with little topographic relief and low slope, while low values are associated with rugged areas of high slope. (f) Optimized cost surface in the Tahoe region for the Comp1.Comp2 surface. Light shading indicates low resistance values, while dark shading indicates areas of high resistance to gene flow. (g) Circuitscape output for the optimized cost surface in the Tahoe region with sampling sites as the focal nodes and the output raster Comp1.Comp2 stack as the resistance surface. Light shading indicates high gene flow current, while dark shading indicates areas of low gene flow.

#### PLUMAS

Our Plumas study site is located in Plumas National Forest (southwest corner of extent: - 121.4079, 39.5963), and is roughly defined by Feather River Canyon in the northwest; tributaries of Lake Oroville in the southwest; Sly Creek Reservoir, La Porte, and Little Grass Valley in the southeast; and the town of Quincy in the northeast (Figure 6), and includes 12 sampling sites (Figure 6a). The only major road in this landscape is a small portion of Highway 70 in the northwest corner of the extent, along Feather River Canyon (Figure 6a). The landscape is largely comprised of forests and woodlands made up of cool temperate forest species— including conifers (e.g., *Pinus* spp., *Sequoia* spp.) and broadleaf evergreen trees (e.g., *Arbutus* spp., *Quercus* spp.), a few large lakes, rivers, and streams. Developed and other human use areas are present along roads and areas of recently disturbed or modified vegetation in the south. Shrub and herb vegetation including Californian scrub (chaparral), desert, and semi-desert areas largely made up of sagebrush scrubland, and open rock vegetation (including non-vascular plant species), among other landscape elements, are present in lower proportions (Figure 6b).

**Figure 6.**
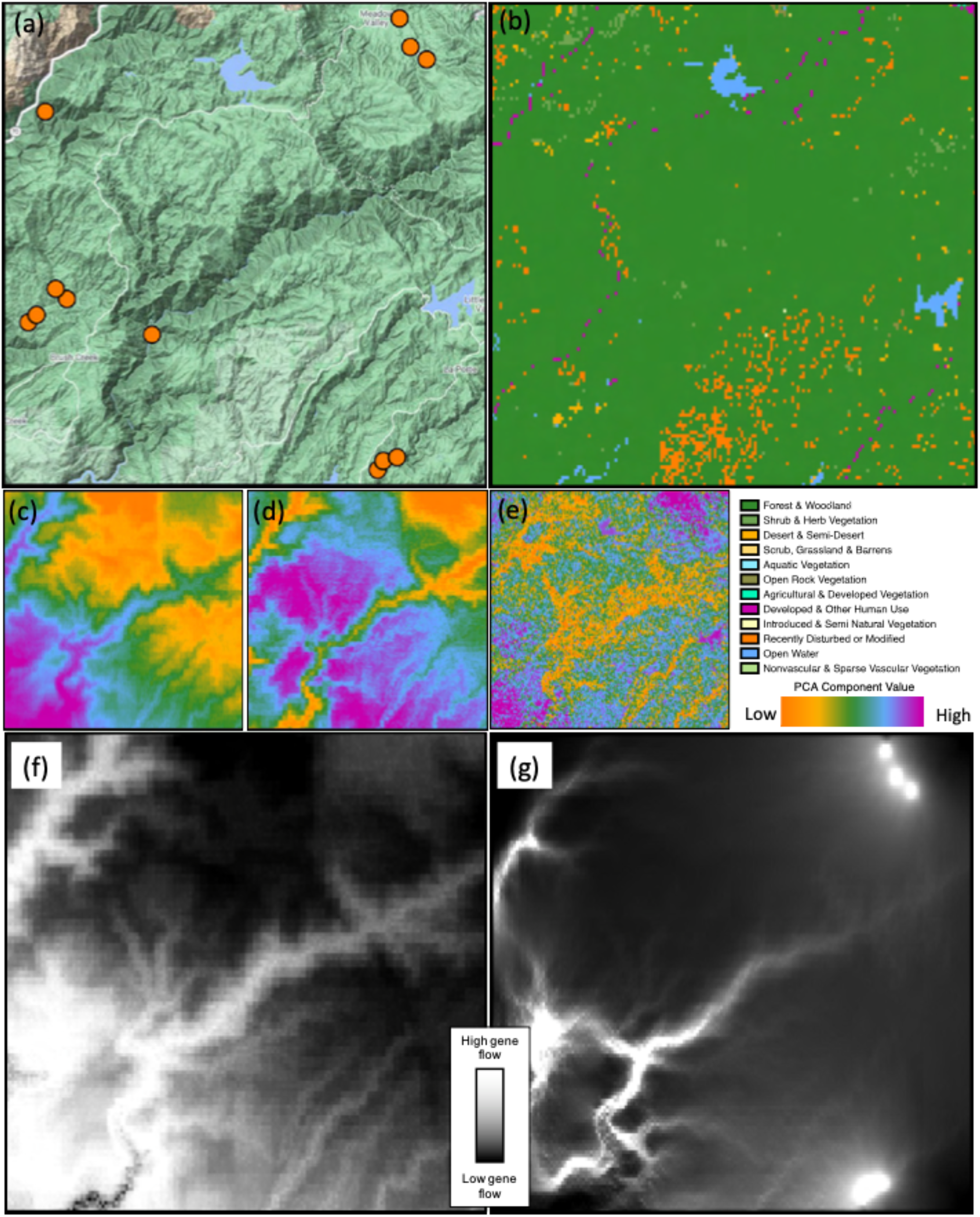
(a) Terrain map of the Plumas region, where orange points indicate sampling locations (*n* =12). Terrain map data © 2021 Google. (b) Vegetation map of the region, with each color representing a distinct vegetation category. (c) Heatmap of PCA component 1, where high values are associated with warmer areas of lower elevation, while low values are associated with wetter areas of higher elevation. (d) Heatmap for PCA component 2, where high values are associated with wetter areas (especially wet season precipitation), while low values are associated with more temperature seasonality. (e) Heatmap for PCA component 3, where high values have little topographic relief and low slope, while low values are associated with rugged areas of high slope. (f) Optimized cost surface in the Plumas region for the Comp1.Comp2.Comp3 surface. Light shading indicates low resistance values, while dark shading indicates areas of high resistance to gene flow. (g) Circuitscape output for the optimized cost surface in the Plumas region with sampling sites as the focal nodes and the output raster Comp1.Comp2.Comp3 as the resistance surface. Light shading indicates high gene flow current, while dark shading indicates areas of low gene flow.

### LANDSCAPE GENETICS ANALYSIS

We used ResistanceGA (Peterman *et al.* 2014; Peterman 2018) for our landscape genetics analyses. ResistanceGA uses a genetic algorithm to simultaneously estimate resistance values from continuous and categorical surfaces based on pairwise genetic data and effective geographic distances. The method is free from subjective *a priori* assumptions of the expected resistance values, and fully explores parameter space. This allows for bias-free estimations of gene flow and migration rates through different areas in the landscape. Here, we largely used default values in the genetic algorithm optimization settings, but selected maximum categorical and continuous values of 500, while evaluating all possible transformations for continuous surface values. We set the maximum number of layer combinations equal to 5, to evaluate models that include all combinations of layers together. We implemented a jackknife resampling procedure of 1000 iterations of 75% of the samples in each analysis and evaluated the results using AICc where all models of delta AICc ≤ 2 of the top model were considered to have substantial support. Finally, we used Circuitscape (version 4; Hall *et al.* 2021) to visualize optimized gene flow across the landscape, with sampling sites as the focal nodes and the output raster as the resistance surface.

## RESULTS

Replicated LG analyses in different areas of the *S. occidentalis* range revealed several points of agreement, and some key differences. We describe results in each area and follow this with a comparison among results.

### SEQUOIA

The layer set with highest support based on AICc values and most common layer set in the jackknife analysis (top model in 86.1% of jackknife analyses) is the composite layer containing PCA component 1, the vegetation layer, and the roads layer (Table S4). High values of PC1 are associated with higher elevation regions with more precipitation, while low values are associated with lower elevation of higher temperatures (Figure S1a, Table S1). The composite layer containing PCA component 2, vegetation, and the roads layer also has substantial support (delta AICc <2; top model in 5.6% of jackknife analyses) (Table S4). For PC2, high values are associated with areas that are more seasonal and experience more dry-season precipitation (Figure S1b, Table S2). These results demonstrate that vegetation, roads, PC1, and PC2 are the primary factors influencing gene flow in this landscape (Table S4).

These results indicate that colder, high-elevation areas that experience high amounts of precipitation (PC1) are barriers to gene flow, and that resistance to gene flow increases rapidly with increasing PC1 values once a threshold is reached (Figure S2). Principal component 2 also substantially influences gene flow, with low values having lowest resistance to gene flow. This suggests highly seasonal areas are also barriers to gene flow (Figure S3). We also find strong support that vegetation influences gene flow in the Sequoia region, where Forest & Woodland and Desert & Semi-Desert habitats present the lowest resistance to gene flow, while Agricultural & Developed Vegetation and Open Water present the strongest barriers to gene flow (Table S5). Finally, we find that roads in this region affect gene flow and appear to be ~5.3x more costly than areas without roads (Table S6).

### SIERRA

The layer set with the highest support based on AICc values and most common layer set in the jackknife analysis (top model in 100% of jackknife analyses) is the composite layer containing PCA components 1, ,2 and 3 (Table S10). High values of PC1 are associated with higher elevation regions with drier seasonal precipitation and cooler temperatures, while low values are associated with lower elevation areas of higher temperatures (Figure S4a, Table S7). For PC2, high values are associated with areas of higher temperature seasonality and higher annual temperature range, while low values are associated with areas of high precipitation (Figure S4b, Table S8). For PC3, high values are associated with areas of high slope and ruggedness, while low values are associated with areas of low slope and ruggedness (Figure S4c, Table S9). These results demonstrate that the environmental axes summarized by PC1, PC2, and PC3 all influence gene flow in this landscape.

These results indicate that colder, high-elevation areas with more precipitation (PC1) are barriers to gene flow, and that resistance to gene flow increases gradually with increasing PC1 values (Figure S5). Principal component 2 also influences gene flow, with intermediate values having lowest resistance to gene flow (Figure S6). This suggests highly seasonal areas and, to a smaller degree, areas of high, wet-season precipitation, are also barriers to gene flow. Additionally, we find support that rugged areas of high slope, as well as areas of very low ruggedness and slope—such as rivers, lakes, and roads—inhibit gene flow (Figure S7). However, we find no support that vegetation or roads spatial layers influence gene flow in the Sierra region in any substantial way (Table S10).

### YOSEMITE

Several layer sets and individual layers are found to influence gene flow in Yosemite based on AICc values (all within delta AICc of 2), such as PC component 2, distance, PC component 1, roads, PC component 3, and the composite layer PC component 2 and roads (Table S14). However, three layers were most often identified as important factors affecting gene flow based on jackknife analyses. These include PC component 2 (top model in 56.5% of jackknife analyses), PC component 1 (top model in 27.2% of jackknife replicates), and geographic distance (top model in 7.8% of jackknife replicates) (Table S14). For PC2, high values are associated with areas of higher wet season precipitation, while low values are associated with areas of higher temperature seasonality and higher annual temperature range, as well as more rugged areas and areas of higher slope (Figure S8b, Table S12). For PC1, high values are associated with higher elevation regions with drier seasonal precipitation and cooler temperatures, while low values are associated with lower elevation areas of higher temperatures (Figure S8a, Table S11).

These results indicate that several factors affect gene flow in the Yosemite region. First, rugged areas of high slope that also have high temperature seasonality and temperature annual range restrict gene flow more so than areas with little topographic relief that experience less variable temperatures (Figure S10). We find further evidence from PC axis 3 that rugged areas of high slope are features that restrict gene flow (Figure S11). We also find evidence that colder, high elevation areas inhibit gene flow, while warmer, lower elevation areas facilitate gene flow (Figure S9). We also observe a substantial effect of isolation by distance—where geographic distance across the landscape is associated with increased genetic differentiation (Table S14). Notably, we find roads strongly inhibit gene flow in the Yosemite region, which are approximately 500x more costly to traverse than areas without roads (Table S15).

### TAHOE

The layer with the highest support based on AICc values and most common layer set in the jackknife analysis (top model in 68.8% of jackknife analyses) is PCA component 2 (Table S19). High values are associated with areas of high temperature seasonality, high temperature annual range, and rugged areas of high slope, while low values are associated with areas that receive more precipitation—especially wet season precipitation (Figure S12b, Table S17). Principal component 3 also has substantial support (delta AICc <2; top model in 22.9% of jackknife analyses) (Table S19). For PC3, high values have little topographic relief and low slope, while low values are associated with rugged areas of high slope (Figure S12c, Table S18).

These results indicate that areas of very high precipitation (Figure S13) and rugged areas of high slope inhibit gene flow (Figure S14). While we find that precipitation does not appear to affect gene flow until it reaches very extreme levels, resistance to gene flow increases continuously with increasing ruggedness and slope (Figures S13, S14). Here we attribute high resistance to gene flow in areas of very low ruggedness to water bodies within this landscape (see discussion for details).

### PLUMAS

Three composite layers that include different combinations of PC axes 1, 2, 3, and roads have the highest support based on AICc values and are all included in the top models in jackknife replicates (Table S23). The layer stack that includes PC1, PC3, and the roads layer has the lowest AICc value and is the most common layer set in the jackknife analysis (top model in 47.2% of jackknife replicates). The layer stack that includes PC1, PC2, and roads also has substantial support (delta AICc <2; top model in 28.2% of jackknife replicates). Likewise, the layer stack that includes PC2, PC3, and roads also has substantial support (delta AICc <2; top model in 24.6% of jackknife replicates) (Table S23). For PC1, high values are associated with warmer areas of lower elevation, while low values are associated with wetter areas of high elevation (Figure S15a, Table S20). For PC2, high values are associated with wetter areas (especially wet-season precipitation), while low values are associated with greater temperature seasonality (Figure S15b, Table S21). Finally, for PC3, high values have little topographic relief and low slope, while low values are associated with rugged areas of high slope (Figure S15c, Table S22).

These results indicate that colder, high elevation areas restrict gene flow, while warm, low elevation areas facilitate gene flow (Figure S16). We also find very wet areas (especially in terms of wet-season precipitation) and areas of high temperature seasonality restrict gene flow (Figure S17). Additionally, areas of high topographic relief and high slope are resistant to gene flow, but areas of low slope and ruggedness—such as rivers, lakes, and roads—inhibit gene flow (Figure S18). Finally, while we find that roads facilitate gene flow on the order of 2.27 times over non-road areas (Table S24), we believe this is an artifact due to only including one small road (Highway 70, Feather River Highway) that occurs in the northwest corner of the extent at the base of Feather River Canyon.

### COMPARISON BETWEEN REGIONS

We identified similarities in barriers to gene flow between several regions. High-elevation, colder areas that often have more dry-season precipitation were identified as barriers to gene flow in most regions (Sequoia, Sierra, Yosemite, and Plumas). Likewise, areas of high ruggedness and slope restricted gene flow significantly in most areas (Sierra, Yosemite, Tahoe and Plumas). High temperature seasonality restricted gene flow in three areas (Sierra, Yosemite, Plumas)—though it should be noted that areas of high temperature seasonality are also most often found at high-elevation regions in our northernmost Plumas study area. Similarly, high precipitation during the wet season was a significant factor limiting gene flow in the three study areas (Sierra, Tahoe, and Plumas). Some barriers to gene flow were less common across study areas, while others appear to be unique. For example, roads were identified as 5.3x more costly in the southernmost study area of Sequoia, while 500x more costly in the Yosemite study area. Areas of high and low gene flow for each area are shown in Figures 2–6, panels f and g. Table 1 presents a comparison of the results between regions.

**Table 1.**
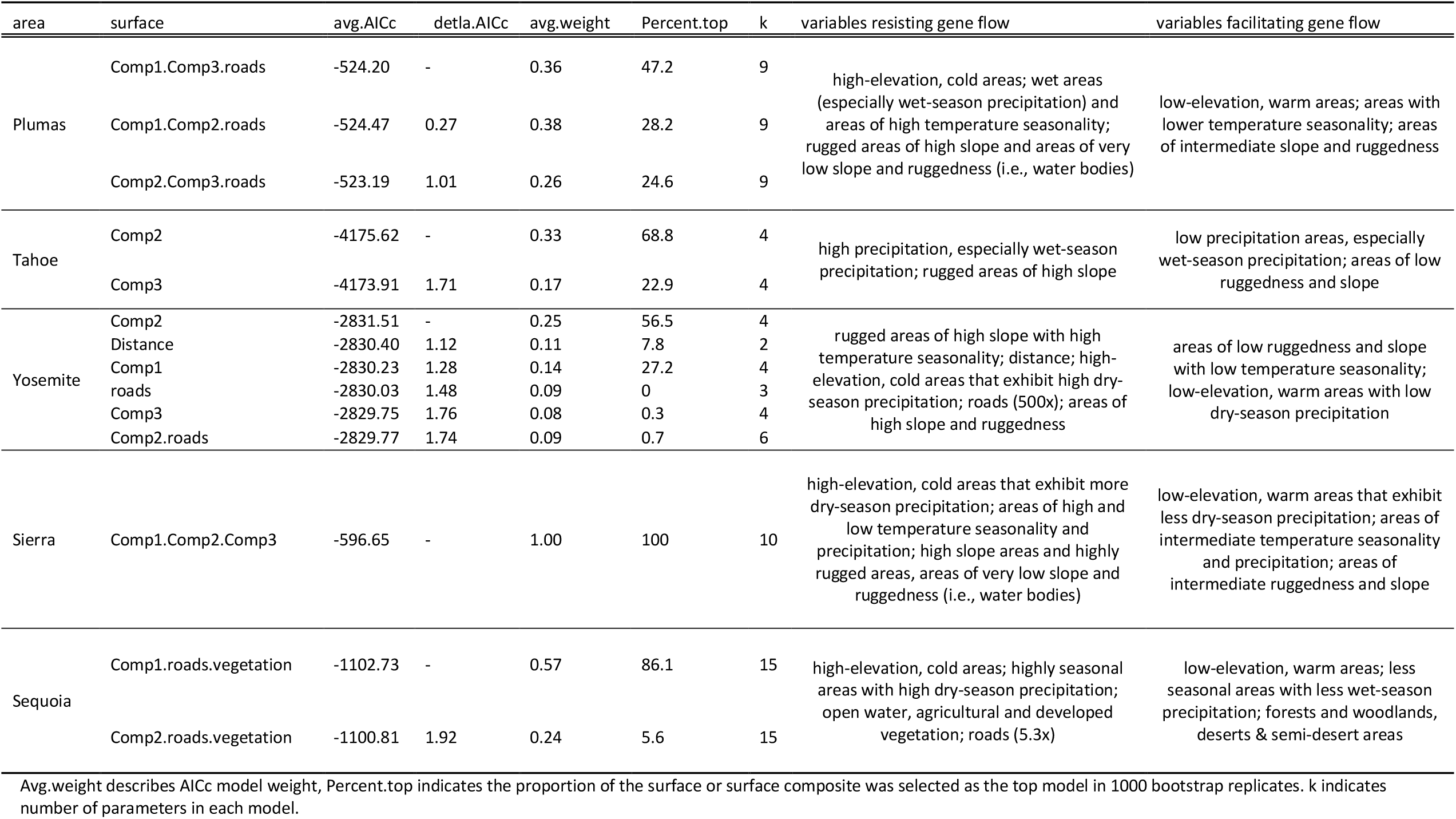
Summary of comparison in barriers and facilitators to gene-flow between the five study areas. The top model (based on AICc) and all models with substantial support (within delta 2 AICc) are shown.

## DISCUSSION

### SIMILARITIES IN RESULTS ACROSS STUDY AREAS

We found a great deal of concordance in the ecological factors that represent major barriers to gene flow for Western Fence lizards (*S. occidentalis*). First, across the study areas Sequoia, Sierra, Yosemite, and Plumas, we found high-elevation colder areas are impediments to gene flow. This result is not surprising, given that Western Fence lizards are not encountered above approximately 3,300 m in elevation (Bell & Price 1996; Stebbins 2003; Leaché *et al.* 2010), where temperatures tends to be colder and the areas experience more dry-season (summer) precipitation. Similarly, we found a strong trend toward topographic features such as ruggedness and slope restricting gene flow, whereas less rugged areas across the landscape enabled gene flow. This was evident in the Sierra, Yosemite, Tahoe, and Plumas study areas. This result is also not surprising, given that areas of high slope, such as cliffs and canyons, often restrict the movement of terrestrial animals across their range (Cushman *et al.* 2006; Richardson 2012; Wang 2012; Bradburd *et al.* 2013; Castillo *et al.* 2014; Van Buskirk & van Rensburg 2020). Temperature seasonality, or standard deviation in annual temperature, also played a major role in restricting dispersal and gene flow across study areas Sierra, Yosemite, and Plumas. This has recently been demonstrated to be an inhibitor of gene flow in the Island Fence lizard *S. occidentalis taylori* (Trumbo *et al.* 2021). Another climate variable that restricts dispersal and gene flow across areas is precipitation—specifically, wet season precipitation— which had a substantial effect in the Sierra, Tahoe, and Plumas areas.

### DIFFERENCES IN RESULTS BETWEEN STUDY AREAS

We also found evidence of idiosyncrasies in the factors impeding dispersal and gene flow between these same areas. For example, the effect of roads on dispersal and gene flow varied greatly among areas. One might expect the effect of roads to vary widely, depending on where roads are located in the landscape, the amount of traffic the road sees, and the number and type of roads in an area (Holderegger & Wagner 2008; Sunnucks & Balkenhol 2015; McCartney-Melstad *et al.* 2018). Our study supports such expectations, with roads being approximately 5x more restrictive to dispersal in the southernmost Sequoia site, to approximately 500x more restrictive to gene flow in the Yosemite study region. These results may be explained by the number of visitors in each area, as the Yosemite region receives more than 4-5 million visitors annually (irma.nps.gov), while Sequoia National Park receives ~1 million visitors annually (irma.nps.gov). The Kern River Canyon area where our study area is receives even fewer visitors annually. Comparing traffic counts at a centrally-located traffic junction within each area (Highway 178 junction with route 155 in Sequoia and Highway 120 junction with Big Oak Flat road) confirms the differences in traffic volumes, though to a smaller degree, with the Yosemite region receiving approximately twice as much traffic (dot.ca.gov/). However, and importantly, while the effect of roads on resistance to dispersal and gene flow can vary widely and be extreme in some cases, roads do not appear to affect long-term genetic structure in Western Fence lizards. Primary and secondary roads make up a very small proportion of the overall environment across the landscape, so their effect is relatively less important, and, in two study areas (Sierra and Tahoe), roads were not found to have a substantial effect. Still, they can represent significant barriers to dispersal and gene flow where they do occur. In one instance, we also found roads were identified as facilitators to dispersal and migration—which is certainly a possibility for this and other species where roads seldom used represent dispersal and migration corridors—but in this case, we believe the result is attributable to the landscape only having one small road in the northwest corner of the extent, at the base of Feather River Canyon. In any case, while we identified this road effect, we did not specifically set out to test the effect of roads in our design and so this result should be interpreted with caution. The analysis of roads on dispersal and genetic connectivity is complex (McCartney-Melstad *et al.* 2018), and confounded by various road types, differences in usage, and other factors—such as underpasses and underpass tunnels that were present in our Tahoe study area, where we did not identify a strong effect of roads on genetic connectivity. In short, analyzing the effect of roads on genetic connectivity is likely to vary across space due to a multitude of factors. Nevertheless, our results reaffirm previous studies that demonstrate the effect of roads as barriers to dispersal and genetic connectivity can depend on the amount of traffic (McCartney-Melstad *et al.* 2018).

Across study areas, while we do find a great deal of consistency in the major barriers to dispersal and genetic connectivity, we also identified some site-specific differences. For example, in the southernmost Sequoia study area, we found substantial support for vegetation type affecting migration and genetic connectivity in this landscape, with the Forest & Woodland habitat representing the lowest cost vegetation type to traverse. Desert & Semi Desert areas seem to be important barriers to gene flow only when they dominate large areas of the landscape. Notably, this study area differs from others in that the extent is not dominated by Forest & Woodland vegetation and/or Scrub, Grassland & Barrens, but also includes substantial Desert & Semi-Desert habitat, Shrub & Herb Vegetation, and Open Rock Vegetation, among others (Figure 2b). In other areas where forest cover is not limiting, or fragmented, the analysis does not identify Forest & Woodland habitat as an important facilitator of gene flow because of the absence of variability in the levels of this categorical predictor variable. This was one of the most important findings in an early study of spatial replication in landscape genetics of black bears, where gene flow is facilitated by forests; however, the analysis failed to detect an effect of forests when they were not variable or fragmented (Short Bull *et al.* 2011). We believe Forest & Woodland, along with Scrub, Grassland & Barrens are generally favorable to dispersal and migration in Western Fence lizards, though not captured by the model in the majority of areas examined here. Instead, elevation, climate, and other topographic features are more variable across the landscape and those differences exert a relatively stronger effect on genetic connectivity across space. In other *Sceloporus* species, however, vegetation types can greatly affect genetic connectivity in different parts of a species range (Clark *et al.* 1999; Branch *et al.* 2003), so it is unlikely vegetation plays no role in dispersal and connectivity, though it seems that vegetation is relatively less important in the Sierra Nevada, where suitable habitat is not limiting.

In another area, Tahoe National Forest, the main finding that colder, higher elevation areas represent barriers to gene flow was not substantially supported. Lizards in this region seem to maintain high levels of genetic connectivity across high elevation areas and are more restricted by areas of high slope and high precipitation. It is unclear if this is a unique attribute of this population, or a limitation in the model. One possible explanation is that the area of highest elevation in this landscape occurs at Donner Pass (~2100 m), which is below the elevation limits of this species (~3300 m), making it difficult to detect the effect of elevation on dispersal and genetic connectivity. This absence of a detected effect is similar to the results found for vegetation across sites, where vegetation type has a substantial effect on gene flow only in areas where ideal habitat is limiting (i.e., the Sequoia region in the south). In the Tahoe region, areas of high elevation that exceed the limits for this species are not present in this landscape, and therefore would not be detected. However, these topographic features are unlikely to be the only factors responsible for this effect, as elevations that do not exceed the range limits of Western Fence lizards are also not found in several other study areas. We believe the unique effect found in Tahoe may result from a combination of factors, including what appears to be a large population in the Donner Summit high-elevation plateau, which apparently maintains high levels of genetic connectivity. As no low-elevation dispersal corridors are present in the northeastern quadrant, dispersal and genetic connectivity across this high-elevation region apparently persists at high enough levels to not be classified as a barrier by the model.

### GENERAL PATTERNS OF LANDSCAPE FACTORS AFFECTING GENETIC CONNECTIVITY

Concordance in landscape genetics results across space is, however, remarkably consistent across study sites when all variables identified as substantially influential to dispersal and genetic connectivity are taken into account. This can be clearly visualized in the optimized cost surface maps and circuitscape output maps parameterized on optimized cost surface data (Figures 2–6 f-g). Taken together, it is clear that, across study areas, canyons are important areas of the landscape for the dispersal and maintenance of genetic connectivity. Examining the PC axis 1 maps along with the optimized cost surface and circuitscape output maps illustrates this point (Figures 2–6 panels b,f,g). This result is clear even in the Sequoia region, where vegetation plays a substantial role in affecting patterns of genetic connectivity (dispersal and genetic connectivity is strong in the Kern River Canyon along Highway 178 in the southwest portion of this region). That is, across study areas, the multivariate axes of environmental variation that affect gene flow may differ in subtle ways. Nevertheless, the environments within these canyons are always substantially implicated as facilitators to dispersal and key features linking populations and maintaining genetic connectivity across these landscapes.

### NOTES ON SPATIAL DATA INTERPRETATION AND MODEL BEHAVIOR

Spatial data are inherently complex and multidimensional, which makes careful assessment of the variables affecting dispersal and genetic connectivity critical both to correct interpretation of the results as well as subsequent interpretations. For example, there may be some vegetation types that are only found in certain environments, making it impossible to separate the effects of a vegetation type from the environmental variables present where they are found. For example, across the Sierra Nevada, the scrub, Grassland & Barrens vegetation class is found almost exclusively at high elevations. This spatial association makes it difficult to distinguish the effect of this vegetation type from the effect of high-elevation, colder areas. In practice, this is not usually an issue, and interpretation of the critical variables affecting dispersal and genetic connectivity should be based on the combination of factors present in any given spatial location, as we have done here. There exists an inherent association between continuous environmental variables and categorical vegetation types that should not be ignored in the interpretation of factors affecting gene flow across landscapes (Peterman & Pope 2020).

The complexity of spatial data layers is also exemplified in the interpretation of the principal components constructed from the 18 temperature, climate, and topographic variables examined here. For each of these axes, we emphasize the interpretation based on the variables that load most strongly with each PC axis. Nevertheless, it should be noted that the remaining environmental and topographic variables are also correlated with these PC axes to varying degrees and should not be ignored. For example, the PCA loadings for component 3 in the Sierra region loads most strongly with topographic ruggedness (0.633) and slope (0.631), and we focus our interpretation of this layer as exemplifying these features. However, precipitation seasonality and mean precipitation are also correlated with Sequoia PC3 to a smaller degree (−0.245 and −0.209, respectively) (Table S3). Environments are experienced on the landscape as the combination of factors present in each area, rather than as isolated variables, and should be interpreted as such (Peterman & Pope 2020). Here, our findings reiterate the perspective that varying degrees of correlation among continuous environmental variables may affect the results, emphasizing the importance of appreciating the multidimensionality of spatial data in their interpretation.

Finally, beyond the issues raised above, continuous topographic and climactic variables may capture unique properties of the environment that are important to recognize. For example, in several study areas (Sierra, Yosemite, Tahoe, Plumas), the ResistanceGA model identified high topographic ruggedness and high slope as areas that inhibit gene flow. This result is both clear and intuitive, as it seems obvious that sheer canyons, for example, can inhibit dispersal and reduce genetic connectivity. However, the model also identifies areas of very low slope and ruggedness as barriers to dispersal and gene flow (Figures S7, S10, S14, and S18). This result is seemingly counterintuitive, given that flat, even landscapes are expected to promote dispersal and gene flow in this species. This result only makes sense in the context of other spatial layers and the features present in those areas. Examining plots of topographic relief in each area along with vegetation plots (Figures 1–5 panels b and e) demonstrates that areas of low topographic slope and ruggedness include water bodies such as rivers and lakes. This occurs because the ResistanceGA model is naïve to the what the landscape variables are and their hypothesized effect on dispersal. Thus, the model correctly identifies water bodies as areas of the landscape that restrict gene flow, but it does so through a PC layer that largely describes microtopographic features. Similarly, in the Plumas region, a small stretch of road occurs only in the northwestern corner of the extent, at the base of Feather River canyon—a dispersal and gene flow corridor in this area. Because the road only occurs in an area that is a high gene flow zone, the ResistanceGA model incorrectly identifies the roads layer as a gene flow corridor. While this may be true in this small area of the landscape, it does not by any means suggest that roads are generally beneficial to dispersal and gene flow for this small lizard. Again, we emphasize that spatial data layers are complex and multidimensional, and a careful consideration of these associations is vital to form sound conclusions about the critical factors affecting dispersal and genetic connectivity across space.

Beyond these issues of data interpretation, we look forward to continued development and refinement of landscape genetics methods and models. While the field of landscape genetics was conceived from simpler methods, such as least-cost paths (LCPs) (Cushman *et al.* 2006; Wang *et al.* 2009; Freedman *et al.* 2010; Van Strien *et al.* 2012; Trumbo *et al.* 2013, 2021; Zancolli *et al.* 2014; Richardson *et al.* 2016) and mantel and partial mantel tests (Andrew *et al.* 2012; Richardson 2012; Hecht *et al.* 2015; Roffler *et al.* 2016; Krohn *et al.* 2019), it is rapidly gaining ground with more sophisticated spatial models that incorporate increasing complexity in the relationships among environmental variables. ResistanceGA represents one area of rapid growth in model development (Peterman 2018; Peterman *et al.* 2019; Winiarski *et al.* 2020). In this study, we implemented what we believe is among the forefront of landscape genetics methods in an isolation by resistance (IBR) framework (Mcrae 2006), and are encouraged by the implementation of jackknife procedures when evaluating model fit. Going forward, we believe the field would benefit greatly from methods that include measures of precision and confidence in the resistance values obtained for continuous and categorial surfaces, which would allow for a quantitative comparison between areas and an understanding of the range of variability in resistance for a given model. Nevertheless, landscape genetics methods as currently implemented provide an important path toward a more nuanced and precise understanding of the critical features of the environment that structure genetic variation (Peterman *et al.* 2014; Peterman 2018; Winiarski *et al.* 2020).

### GENERAL CONCLUSIONS

We predicted that if genetic connectivity is a species-level process, then the barriers to gene flow will be similar across the range of a species. That is, the same landscape variables will show up as important across areas. On the other hand, if barriers and facilitators to dispersal and genetic connectivity are different across the range of the species, then this could mean several things: genetic connectivity may be a population-level process, with different populations within a species experiencing the landscape differently. Population-scale differences may be due to a variety of factors, such as unique behavioral, morphological, or physiological adaptations within each population, or a property of the landscape where habitat types, climate, topography, and distance exert different effects in different areas. Another distinct possibility is that landscapes simply cannot be replicated, since each is unique and may harbor complex underlying structures (such as multilevel interactions between vegetation types, elevation, climate, and geographic distance, interactions with conspecifics, predator-prey relationships, etc.) that make genetic connectivity a unique function of local landscape characteristics in space and time. In a recent review of seascape genetics studies, temperature, oceanography and geography largely dominate as important factors affecting genetic connectivity, idiosyncrasies were also found between species and in different areas (Selkoe *et al.* 2016).

Overall, our results are encouraging for landscape genetics studies. We found that at least for Western Fence lizards across the Sierra Nevada, results are highly congruent, and the same variables often show up as important mediators to dispersal and gene flow in different areas of their range. However, each of the areas also has unique attributes that impose constraints on gene flow. As such, genetic connectivity seems to be affected by similar features of the landscape, and influenced by site-specific factors as well. Taken together, we find that across areas, low-elevation canyons are important dispersal corridors for this species that maintain genetic connectivity across the landscape, which is a result consistent across study areas.

Here we have focused our study on a broadly distributed, common, widespread, generalist ectotherm across a landscape that shares many characteristics in terms of climate, topography, and vegetation across the study areas. We believe an important next step to build upon this work and gain a broader understanding of how the environment influences dispersal and genetic connectivity in nature is to test broad-scale patterns across species. Specialists, for example, might be expected to show results that are even more consistent across space and exhibit less variability within distinct areas. With enough species studied, we can begin to build a theoretical framework to understand how species traits can be used to predict what aspects of the environment are important for dispersal and genetic connectivity, and how consistent these patterns are across a species’ range. Ultimately, this approach may provide valuable insights into how genetic diversity in species is affected and maintained by the environments in which they live and how their genetic structures change over time.

## ACKNOWLEDGEMENTS

We are grateful to A. Leaché and S. Bouzid for contributing samples they had collected in Yosemite National Park. Special thanks to G. Pauly for advice on collecting lizards and for contributing materials and supplies needed to preserve museum specimens. We also thank A.B. Musgrove for her help in the field, A.J. Barley who provided guidance with ddRAD lab work and genomic data processing and M. Hadfield and the Kewalo Marine Lab for providing access to Pippin Prep equipment for DNA size-selection. We are also grateful to M. Kakimoto for her keen attention to detail as an editor and first reader. I am deeply indebted to my PhD committee members: Rob Toonen, Floyd Reed, Amber Wright, and Daniel Rubinoff for their intellectual guidance and expert advice. The technical support and advanced computing resources from the University of Hawaii Information Technology Services – Cyberinfrastructure are gratefully acknowledged. This study was funded by the Theodore Roosevelt Memorial Grant from the American Museum of Natural History, the RCUH Fellowship from the University of Hawaii, the Watson T. Yoshimoto Fellowship from University of Hawaii Ecology, Evolution and Conservation Biology program, the Systematic Research Fund from the Linnean Society of London and the Systematics Association, the Jones-Lovich Research Grant in Southwestern Herpetology from the Society for the Study of Amphibians and Reptiles, a Grant-in-Aid of Research from the Society of Integrative and Comparative Biology, a Grant-in-Aid or Research from Sigma Xi, a University of Hawaii GSO grant, and an NSF grant DEB 1754350 awarded to R.C. Thomson. Euthanasia methods followed those in protocol 16-2384, approved by the UH Institutional Animal Care and Use Committee. All research was conducted under Scientific Collecting Permit SC-13472 issued to V. Wishingrad by the California Department of Fish and Wildlife. We acknowledge the Indigenous lands on which this study took place, including those of the Kawaiisu, Numu, Tübatulabal, Yokuts, Mono, Monache, Newe, Chukchansi, Me-Wuk, Washoe, Nisenan, Kojomk’aki, Maidu, Yana, Atsugewi, Achumawi, and Modoc.

