## Supplemental materials for "Testing concordance and conflict in spatial replication of landscape genetics inferences"

**Supplemental Material for “Testing concordance and conflict in spatial replication of landscape genetics inferences”**

### Sequoia

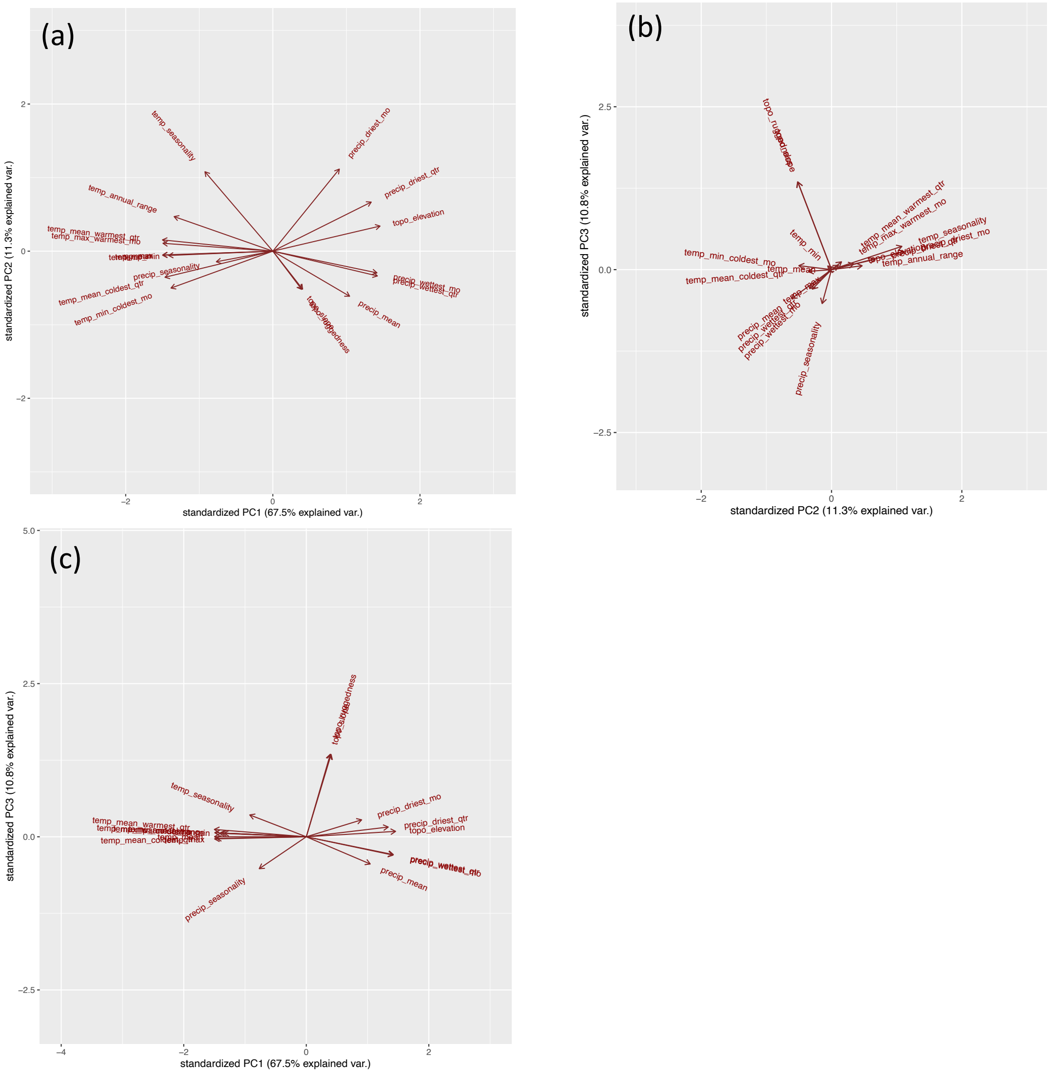


Figure S1. PCA plots of temperature, precipitation, and topography in the Sequoia region. Panel (a) shows component 1 vs component 2; panel (b) shows component 2 vs component 3; and panel (c) shows component 1 vs component 3.

Table S1. PCA loadings for component 1 in the Sequoia region, which explains 67.5% of the variation in the environment.

| Surface | PC1 |
|---|---|
| topo_elevation | 0.273 |
| precip_wettest_qtr | 0.266 |
| precip_wettest_mo | 0.265 |
| precip_driest_qtr | 0.251 |
| precip_mean | 0.196 |
| precip_driest_mo | 0.169 |
| topo_slope | 0.076 |
| topo_ruggedness | 0.073 |
| precip_seasonality | -0.144 |
| temp_seasonality | -0.173 |
| temp_annual_range | -0.252 |
| temp_min_coldest_mo | -0.260 |
| temp_min | -0.266 |
| temp_mean_coldest_qtr | -0.275 |
| temp_max_warmest_mo | -0.280 |
| temp_max | -0.281 |
| temp_mean | -0.281 |
| temp_mean_warmest_qtr | -0.281 |

Table S2. PCA loadings for component 2 in the Sequoia region, which explains 11.3% of the variation in the environment.

| Surface | PC2 |
|---|---|
| precip_driest_mo | 0.512 |
| temp_seasonality | 0.496 |
| precip_driest_qtr | 0.306 |
| temp_annual_range | 0.216 |
| topo_elevation | 0.156 |
| temp_mean_warmest_qtr | 0.070 |
| temp_max_warmest_mo | 0.049 |
| temp_max | -0.024 |
| temp_mean | -0.025 |
| temp_min | -0.027 |
| precip_seasonality | -0.066 |
| precip_wettest_mo | -0.135 |
| precip_wettest_qtr | -0.155 |
| temp_mean_coldest_qtr | -0.163 |
| temp_min_coldest_mo | -0.230 |
| topo_slope | -0.237 |
| topo_ruggedness | -0.238 |
| precip_mean | -0.281 |

Table S3. PCA loadings for component 3 in the Sequoia region, which explains 10.8% of the variation in the environment.

| Surface | PC3 |
|---|---|
| topo_ruggedness | 0.633 |
| topo_slope | 0.631 |
| temp_seasonality | 0.168 |
| precip_driest_mo | 0.130 |
| precip_driest_qtr | 0.075 |
| temp_mean_warmest_qtr | 0.056 |
| topo_elevation | 0.042 |
| temp_max_warmest_mo | 0.031 |
| temp_min_coldest_mo | 0.029 |
| temp_annual_range | 0.028 |
| temp_min | 0.027 |
| temp_mean | 0.001 |
| temp_mean_coldest_qtr | -0.013 |
| temp_max | -0.016 |
| precip_wettest_qtr | -0.137 |
| precip_wettest_mo | -0.141 |
| precip_mean | -0.209 |
| precip_seasonality | -0.245 |

Table S4. Jackknife results for landscape genetics analysis in the Sequoia region.

| surface | avg.AIC | avg.AICc | avg.weight | avg.rank | avg.R2m | avg.LL | n | Percent.top | k |
|---|---|---|---|---|---|---|---|---|---|
| Comp1.roads.vegetation | -862.7321179 | -1102.732118 | 0.566396967 | 1.146 | 0.265991539 | 435.366059 | 861 | 86.1 | 15 |
| Comp2.roads.vegetation | -860.8139542 | -1100.813954 | 0.242830167 | 2.182 | 0.295282604 | 434.4069771 | 56 | 5.6 | 15 |
| Comp3.roads.vegetation | -860.1271282 | -1100.127128 | 0.190772867 | 2.672 | 0.233862491 | 434.0635641 | 83 | 8.3 | 15 |
| Comp1.Comp3.vegetation | -862.4021933 | -1043.735527 | 8.79E-14 | 4.113 | 0.212482099 | 435.2010966 | 0 | 0 | 16 |
| Comp2.Comp3.vegetation | -861.0365845 | -1042.369918 | 4.63E-14 | 5.21 | 0.238190707 | 434.5182922 | 0 | 0 | 16 |
| Comp1.Comp2.vegetation | -860.2123078 | -1041.545641 | 3.80E-14 | 5.677 | 0.235895106 | 434.1061539 | 0 | 0 | 16 |
| Comp1.Comp3.roads.vegetation | -863.7120604 | -1000.51206 | 4.43E-23 | 7.304 | 0.263200167 | 435.8560302 | 0 | 0 | 18 |
| Comp1.Comp2.roads.vegetation | -862.3185222 | -999.1185222 | 2.10E-23 | 7.897 | 0.304604171 | 435.1592611 | 0 | 0 | 18 |
| Comp2.Comp3.roads.vegetation | -861.1535544 | -997.9535544 | 1.13E-23 | 8.799 | 0.272030167 | 434.5767772 | 0 | 0 | 18 |
| Comp1.Comp2.Comp3.vegetation | -859.5471627 | -986.2138294 | 3.59E-26 | 10.003 | 0.200476579 | 433.7735813 | 0 | 0 | 19 |
| Comp1.Comp2.Comp3.roads.vegetation | -861.5744414 | -977.0744414 | 3.28E-28 | 10.997 | 0.232049192 | 434.7872207 | 0 | 0 | 21 |
| Comp1 | -858.8714486 | -854.4270042 | 1.58E-54 | 12.73 | 0.223458198 | 433.4357243 | 0 | 0 | 4 |
| Comp2 | -857.8785903 | -853.4341459 | 1.40E-54 | 13.206 | 0.169710393 | 432.9392952 | 0 | 0 | 4 |
| Distance | -852.9606269 | -851.8697178 | 7.05E-55 | 14.017 | 0.079271684 | 430.4803135 | 0 | 0 | 2 |
| Comp3 | -854.6308491 | -850.1864046 | 4.52E-55 | 15.325 | 0.161270504 | 431.3154245 | 0 | 0 | 4 |
| roads | -853.0269671 | -850.6269671 | 3.81E-55 | 15.484 | 0.095690014 | 430.5134835 | 0 | 0 | 3 |
| Comp1.roads | -859.3788203 | -847.3788203 | 4.32E-56 | 16.951 | 0.228339902 | 433.6894101 | 0 | 0 | 6 |
| Comp2.roads | -858.1284072 | -846.1284072 | 3.52E-56 | 17.617 | 0.177385907 | 433.0642036 | 0 | 0 | 6 |
| Comp3.roads | -854.7589483 | -842.7589483 | 9.90E-57 | 19.261 | 0.167016148 | 431.3794741 | 0 | 0 | 6 |
| Comp1.Comp2 | -858.7222548 | -840.0555881 | 1.09E-57 | 20.49 | 0.178764887 | 433.3611274 | 0 | 0 | 7 |
| Comp2.Comp3 | -858.4175474 | -839.7508807 | 1.29E-57 | 20.715 | 0.156768276 | 433.2087737 | 0 | 0 | 7 |
| Comp1.Comp3 | -858.0305162 | -839.3638496 | 5.73E-58 | 21.204 | 0.177089878 | 433.0152581 | 0 | 0 | 7 |
| Comp2.Comp3.roads | -858.6044374 | -813.6044374 | 2.54E-63 | 23.747 | 0.168517995 | 433.3022187 | 0 | 0 | 9 |
| Comp1.Comp3.roads | -858.9207307 | -813.9207307 | 1.70E-63 | 23.987 | 0.20395525 | 433.4603654 | 0 | 0 | 9 |
| Comp1.Comp2.roads | -858.287465 | -813.287465 | 1.80E-63 | 24.266 | 0.202890889 | 433.1437325 | 0 | 0 | 9 |
| Comp1.Comp2.Comp3 | -859.2118543 | -785.878521 | 1.45E-69 | 26.139 | 0.180670591 | 433.6059272 | 0 | 0 | 10 |
| vegetation | -855.6312271 | -782.2978938 | 3.76E-70 | 26.861 | 0.120089999 | 431.8156136 | 0 | 0 | 10 |

| | | | | | | | | | |
|---|---|---|---|---|---|---|---|---|---|
| Comp1.Comp2.Comp3.roads | -857.5476069 | -545.5476069 | 1.12E-121 | 28.491 | 0.18805759 | 432.7738035 | 0 | 0 | 12 |
| roads.vegetation | -859.6491624 | -547.6491624 | 8.48E-121 | 28.509 | 0.189712008 | 433.8245812 | 0 | 0 | 12 |
| Comp1.vegetation | -862.0441006 | Inf | 0 | 30 | 0.249861672 | 435.0220503 | 0 | 0 | 13 |
| Comp2.vegetation | -860.2730616 | Inf | 0 | 30 | 0.313820003 | 434.1365308 | 0 | 0 | 13 |
| Comp3.vegetation | -860.1916433 | Inf | 0 | 30 | 0.211713486 | 434.0958217 | 0 | 0 | 13 |

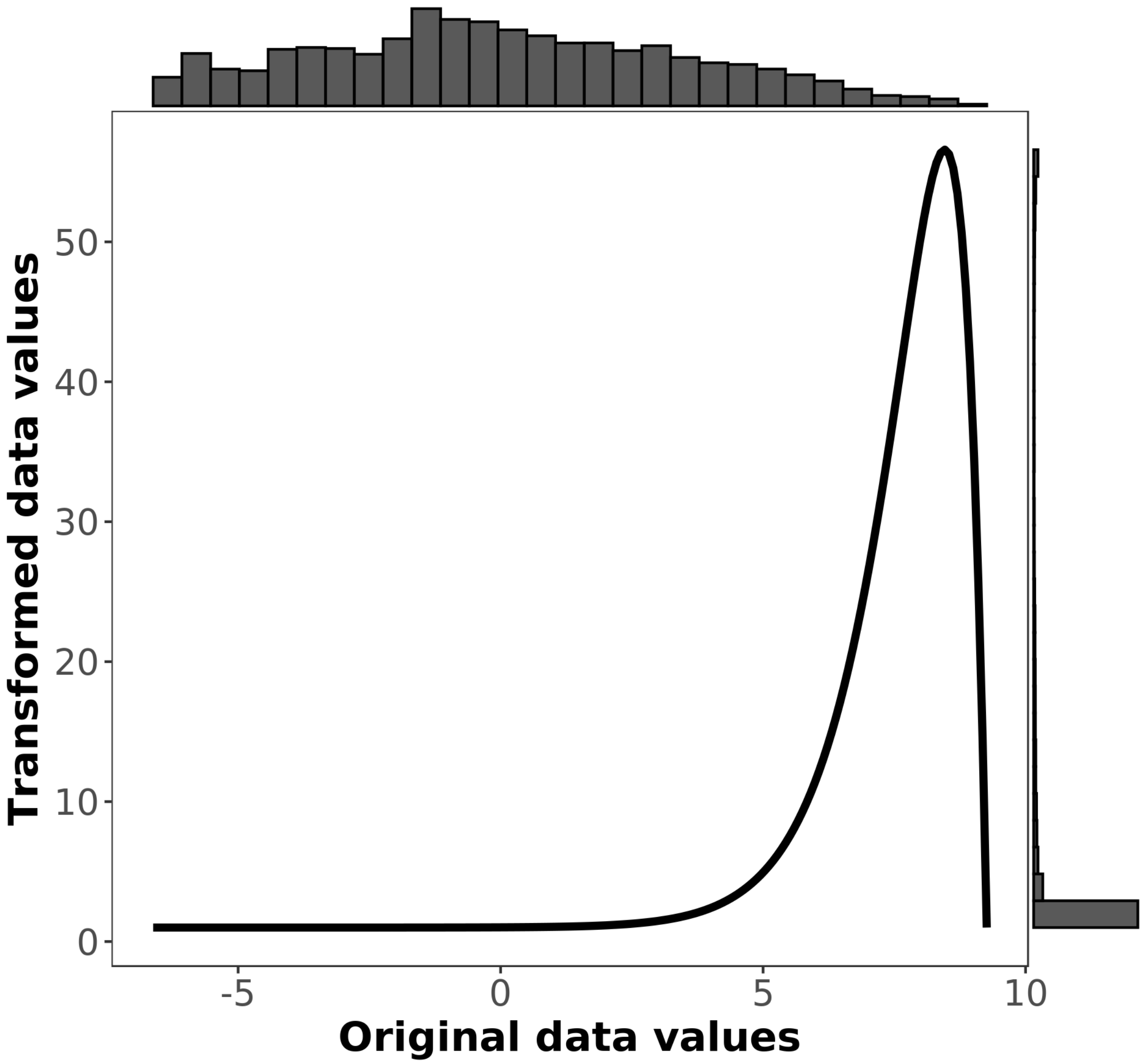


Figure S2. Resistance curve for PC axis 1, which explains 67.5% of the variation in the environment. High values are associated with higher elevation regions with more precipitation and cooler temperatures, while low values are associated with lower elevation areas of higher temperature. Transformed data values indicate the estimated resistance for a given value along the PC axis.

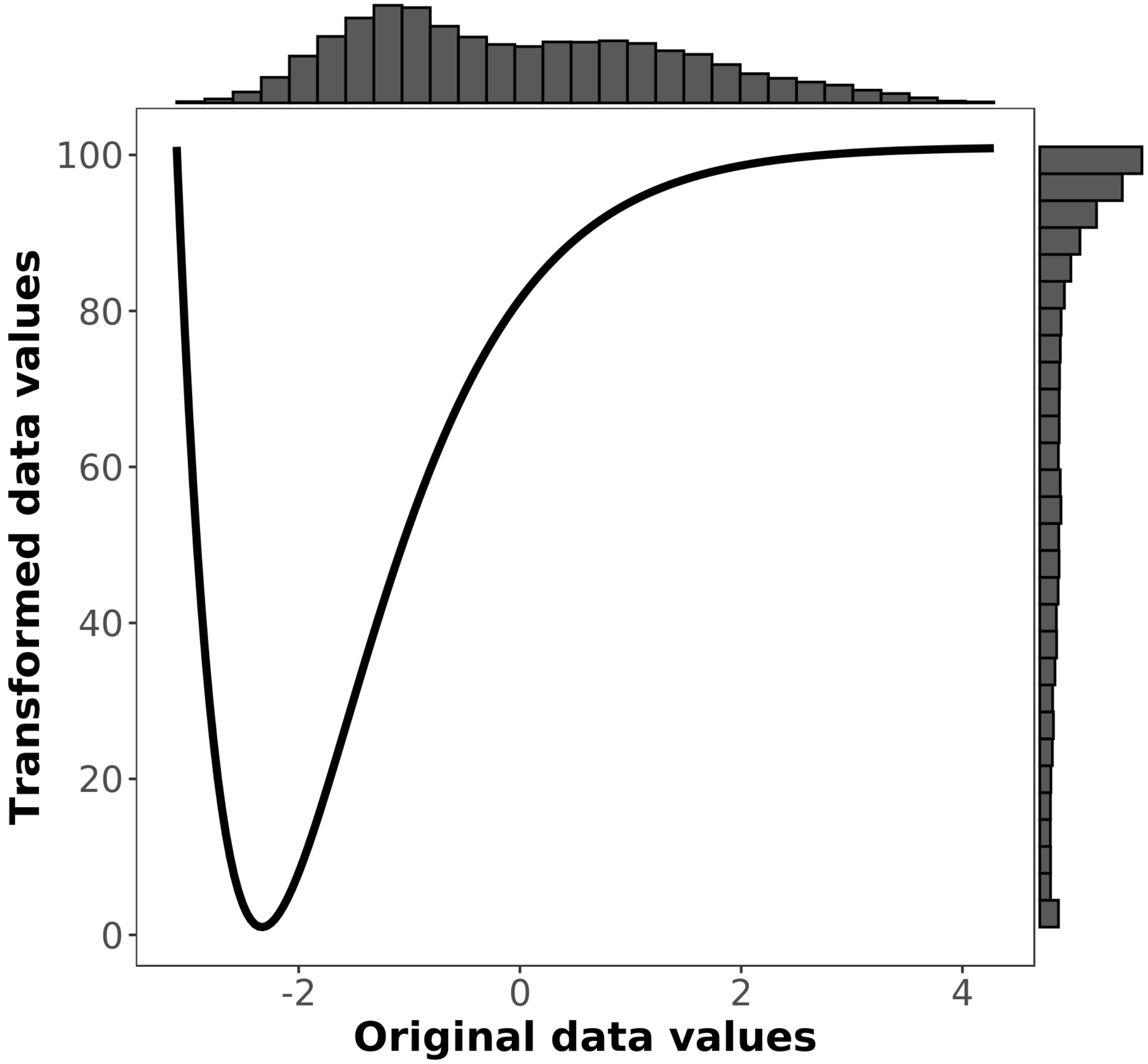


Figure S3. Resistance curve for PC axis 2, which explains 11.3% of the variation in the environment. High values are associated with areas that are more seasonal and experience more dry-season precipitation. Transformed data values indicate the estimated resistance for a given value along the PC axis.

Table S5. Vegetation resistance values in the Sequoia region.

| Vegetation category | Resistance value |
|---|---|
| Forest & Woodland | 1 |
| Desert & Semi-Desert | 2.18 |
| Shrub & Herb Vegetation | 8.06 |
| Recently Disturbed or Modified | 9.57 |
| Open Rock Vegetation | 25.91 |
| Scrub, Grassland & Barrens | 156.29 |
| Developed & Other Human Use | 242.55 |
| Open Water | 269.45 |
| Agricultural & Developed Vegetation | 376.66 |

Table S6. Resistance value for roads in the Sequoia region.

| Feature | Resistance value |
|---|---|
| No road | 1 |
| Road | 5.33 |

### Sierra

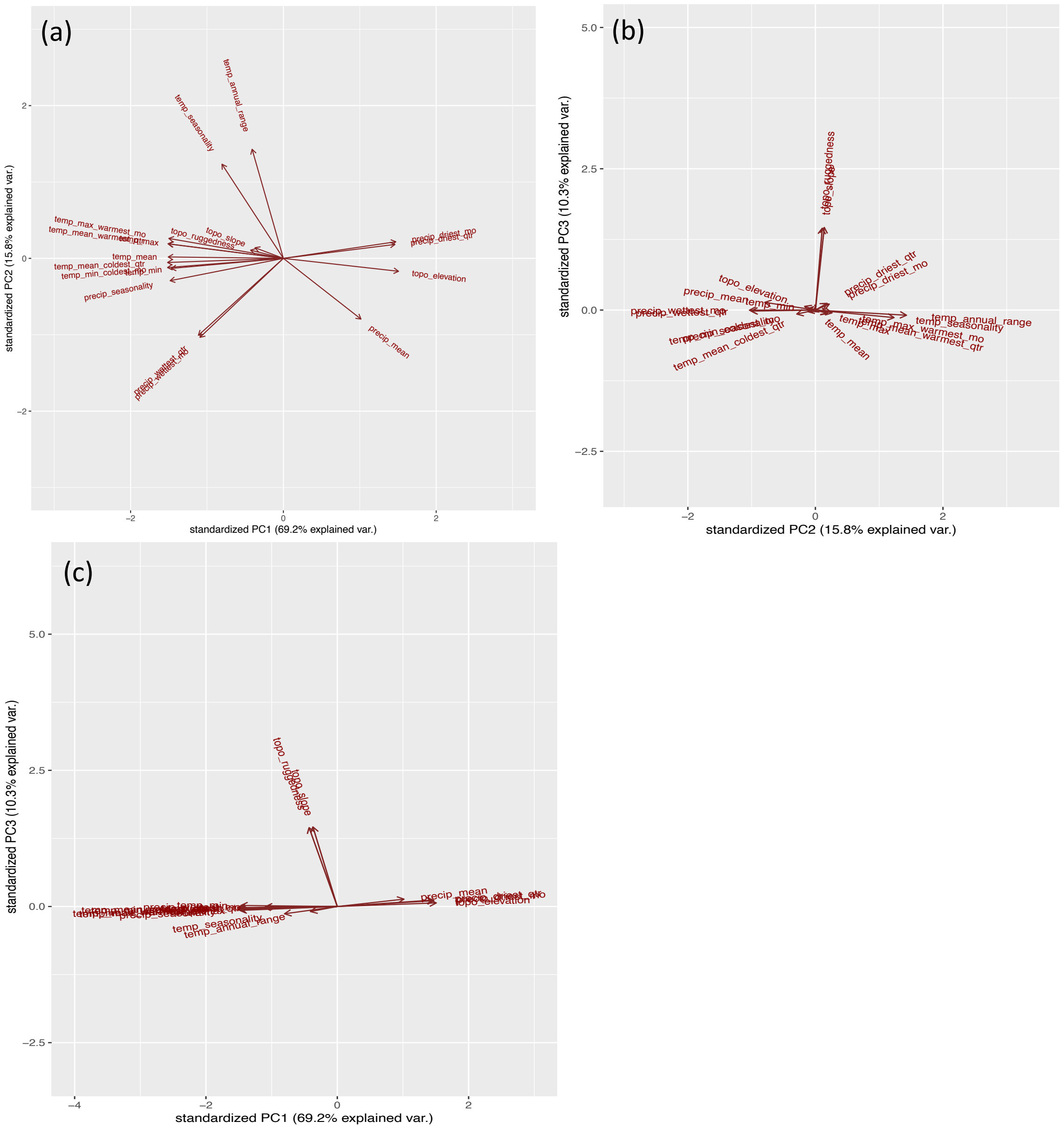


Figure S4. PCA plots of temperature, precipitation, and topography in the Sierra region. Panel (a) shows component 1 vs component 2; panel (b) shows component 2 vs component 3; and panel (c) shows component 1 vs component 3.

Table S7. PCA loadings for component 1 in the Sierra region, which explains 69.2% of the variation in the environment.

| Surface | PC1 |
|---|---|
| topo_elevation | 0.2792 |
| precip_driest_mo | 0.2727 |
| precip_driest_qtr | 0.2705 |
| precip_mean | 0.1891 |
| topo_slope | -0.0676 |
| temp_annual_range | -0.0780 |
| topo_ruggedness | -0.0782 |
| temp_seasonality | -0.1504 |
| precip_wettest_mo | -0.2004 |
| precip_wettest_qtr | -0.2044 |
| temp_min | -0.2718 |
| precip_seasonality | -0.2744 |
| temp_max_warmest_mo | -0.2778 |
| temp_max | -0.2780 |
| temp_mean | -0.2790 |
| temp_mean_warmest_qtr | -0.2798 |
| temp_mean_coldest_qtr | -0.2807 |
| temp_min_coldest_mo | -0.2808 |

Table S8. PCA loadings for component 2 in the Sierra region, which explains 15.8% of the variation in the environment.

| Surface | PC2 |
|---|---|
| temp_annual_range | 0.5509 |
| temp_seasonality | 0.4757 |
| temp_max_warmest_mo | 0.1000 |
| precip_driest_mo | 0.0834 |
| temp_mean_warmest_qtr | 0.0737 |
| temp_max | 0.0733 |
| precip_driest_qtr | 0.0698 |
| topo_slope | 0.0440 |
| topo_ruggedness | 0.0312 |
| temp_mean | 0.0069 |
| temp_mean_coldest_qtr | -0.0210 |
| temp_min_coldest_mo | -0.0465 |
| temp_min | -0.0518 |
| topo_elevation | -0.0658 |
| precip_seasonality | -0.1139 |
| precip_mean | -0.3075 |
| precip_wettest_qtr | -0.3930 |
| precip_wettest_mo | -0.4038 |

Table S9. PCA loadings for component 3 in the Sierra region, which explains 10.3% of the variation in the environment.

| Surface | PC3 |
|---|---|
| topo_slope | 0.7037 |
| topo_ruggedness | 0.6972 |
| precip_mean | 0.0612 |
| precip_driest_qtr | 0.0573 |
| precip_driest_mo | 0.0516 |
| topo_elevation | 0.0306 |
| temp_min | 0.0084 |
| precip_wettest_mo | -0.0053 |
| temp_mean | -0.0088 |
| temp_mean_coldest_qtr | -0.0121 |
| precip_wettest_qtr | -0.0129 |
| temp_min_coldest_mo | -0.0146 |
| temp_mean_warmest_qtr | -0.0220 |
| temp_max_warmest_mo | -0.0223 |
| temp_max | -0.0281 |
| temp_annual_range | -0.0344 |
| precip_seasonality | -0.0351 |
| temp_seasonality | -0.0568 |

Table S10. Jackknife results for landscape genetics analysis in the Sierra region.

| surface | avg.AIC | avg.AICc | avg.weight | avg.rank | avg.R2m | avg.LL | n | Percent.top | k |
|---|---|---|---|---|---|---|---|---|---|
| Comp1.Comp2.Comp3 | -376.6516945 | -596.6516945 | 1 | 1 | 0.043444479 | 192.3258473 | 1000 | 100 | 10 |
| roads.vegetation | -376.8414041 | -508.8414041 | 9.54E-20 | 2 | 0.068111197 | 192.4207021 | 0 | 0 | 11 |
| Comp1.Comp2.Comp3.roads | -376.9598399 | -480.9598399 | 8.14E-26 | 3.699 | 0.070328524 | 192.4799199 | 0 | 0 | 12 |
| Comp3.vegetation | -376.5394892 | -480.5394892 | 6.76E-26 | 4.29 | 0.023423738 | 192.2697446 | 0 | 0 | 12 |
| Comp2.vegetation | -376.502637 | -480.502637 | 6.40E-26 | 5.003 | 0.054700394 | 192.2513185 | 0 | 0 | 12 |
| Comp1.vegetation | -376.4728375 | -480.4728375 | 6.53E-26 | 5.008 | 0.023398977 | 192.2364188 | 0 | 0 | 12 |
| Comp2.roads.vegetation | -376.610506 | -460.610506 | 3.07E-30 | 7.749 | 0.093169063 | 192.305253 | 0 | 0 | 14 |
| Comp1.roads.vegetation | -376.4722229 | -460.4722229 | 2.96E-30 | 8.148 | 0.06829649 | 192.2361114 | 0 | 0 | 14 |
| Comp3.roads.vegetation | -376.4358725 | -460.4358725 | 2.94E-30 | 8.183 | 0.08687099 | 192.2179363 | 0 | 0 | 14 |
| Comp1.Comp2.vegetation | -376.613976 | -456.613976 | 4.01E-31 | 10.514 | 0.038651115 | 192.306988 | 0 | 0 | 15 |
| Comp1.Comp3.vegetation | -376.3914555 | -456.3914555 | 3.68E-31 | 11.187 | 0.031235524 | 192.1957277 | 0 | 0 | 15 |
| Comp2.Comp3.vegetation | -376.470134 | -456.470134 | 3.79E-31 | 11.236 | 0.051285669 | 192.235067 | 0 | 0 | 15 |
| Comp1.Comp2.Comp3.roads.vegetation | -376.8114631 | -453.1750995 | 7.04E-32 | 14.402 | 0.049018691 | 192.4057315 | 0 | 0 | 20 |
| Comp1.Comp2.roads.vegetation | -376.608296 | -453.108296 | 7.36E-32 | 14.5 | 0.056790474 | 192.304148 | 0 | 0 | 17 |
| Comp2.Comp3.roads.vegetation | -376.5011239 | -453.0011239 | 6.76E-32 | 14.813 | 0.029425662 | 192.250562 | 0 | 0 | 17 |
| Comp1.Comp3.roads.vegetation | -376.4046622 | -452.9046622 | 6.66E-32 | 15.044 | 0.023580009 | 192.2023311 | 0 | 0 | 17 |
| Comp1.Comp2.Comp3.vegetation | -376.5328169 | -452.5328169 | 5.18E-32 | 16.224 | 0.052632909 | 192.2664085 | 0 | 0 | 18 |
| Distance | -375.9120208 | -374.1977351 | 5.29E-49 | 18.005 | 0.019924312 | 191.9560104 | 0 | 0 | 2 |
| roads | -376.1268797 | -372.1268797 | 1.94E-49 | 19.028 | 0.060508424 | 192.0634398 | 0 | 0 | 3 |
| Comp1 | -376.4053295 | -368.4053295 | 2.94E-50 | 20.883 | 0.046825206 | 192.2026647 | 0 | 0 | 4 |
| Comp2 | -376.1984633 | -368.1984633 | 2.61E-50 | 20.905 | 0.034452869 | 192.0992317 | 0 | 0 | 4 |
| Comp3 | -376.0028581 | -368.0028581 | 2.76E-50 | 21.179 | 0.046958907 | 192.001429 | 0 | 0 | 4 |
| Comp1.roads | -376.6199923 | -348.6199923 | 1.46E-54 | 23.809 | 0.08224502 | 192.3099962 | 0 | 0 | 6 |
| Comp2.roads | -376.3822102 | -348.3822102 | 1.37E-54 | 23.861 | 0.065923564 | 192.1911051 | 0 | 0 | 6 |
| Comp3.roads | -376.1154807 | -348.1154807 | 1.35E-54 | 24.33 | 0.076048833 | 192.0577404 | 0 | 0 | 6 |
| Comp1.Comp2 | -376.782817 | -320.782817 | 1.27E-60 | 26.503 | 0.041694333 | 192.3914085 | 0 | 0 | 7 |
| Comp2.Comp3 | -376.1744712 | -320.1744712 | 1.01E-60 | 27.241 | 0.031730296 | 192.0872356 | 0 | 0 | 7 |

| | | | | | | | | | |
|---|---|---|---|---|---|---|---|---|---|
| Comp1.Comp3 | -376.2498445 | -320.2498445 | 1.04E-60 | 27.256 | 0.033342586 | 192.1249222 | 0 | 0 | 7 |
| vegetation | -376.5364396 | Inf | 0 | 29 | 0.029459284 | 192.2682198 | 0 | 0 | 9 |
| Comp1.Comp2.roads | -376.8961552 | Inf | 0 | 29 | 0.070195972 | 192.4480776 | 0 | 0 | 9 |
| Comp1.Comp3.roads | -376.618694 | Inf | 0 | 29 | 0.078443992 | 192.309347 | 0 | 0 | 9 |
| Comp2.Comp3.roads | -376.3733128 | Inf | 0 | 29 | 0.055207605 | 192.1866564 | 0 | 0 | 9 |

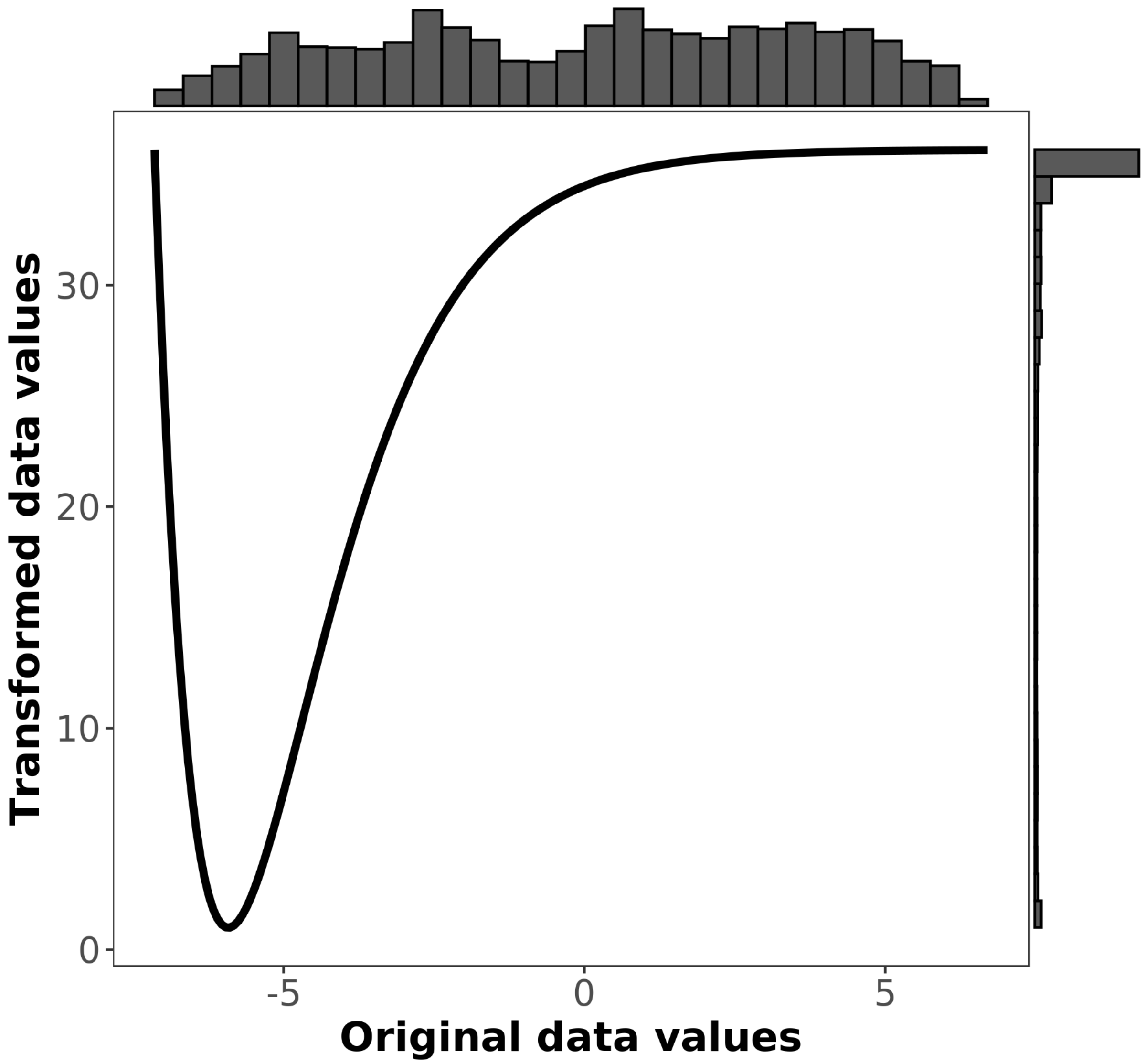


Figure S5. Resistance curve for PC axis 1, which explains 69.2% of the variation in the environment. High values are associated with higher elevation regions with more dry season precipitation and cooler temperatures, while low values are associated with lower elevation areas of higher temperature. Transformed data values indicate the estimated resistance for a given value along the PC axis.

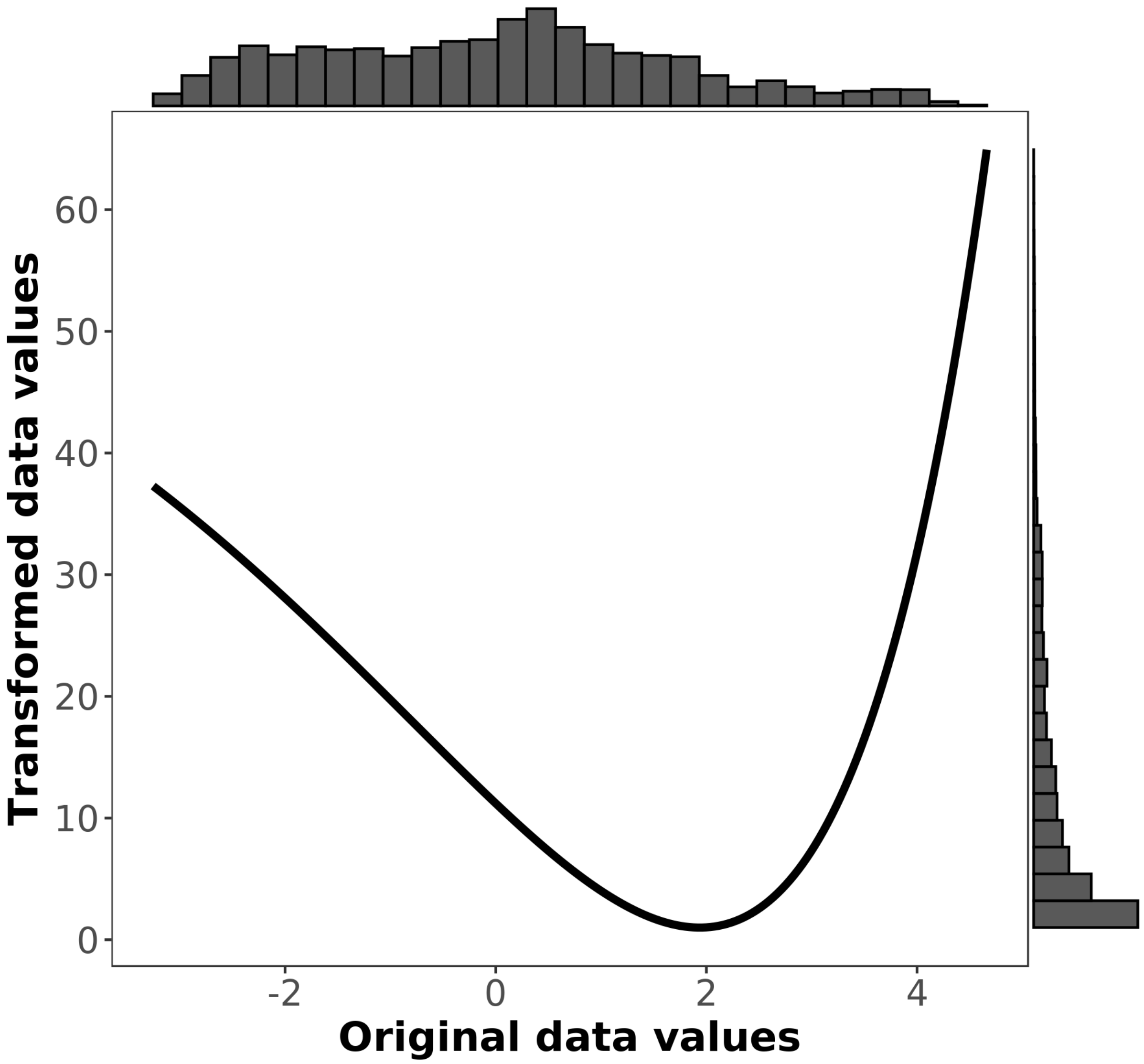


Figure S6. Resistance curve for PC axis 2, which explains 15.8% of the variation in the environment. High values are associated areas of higher temperature seasonality and higher annual temperature range, while low values are associated with areas of high precipitation, especially wet season precipitation. Transformed data values indicate the estimated resistance for a given value along the PC axis.

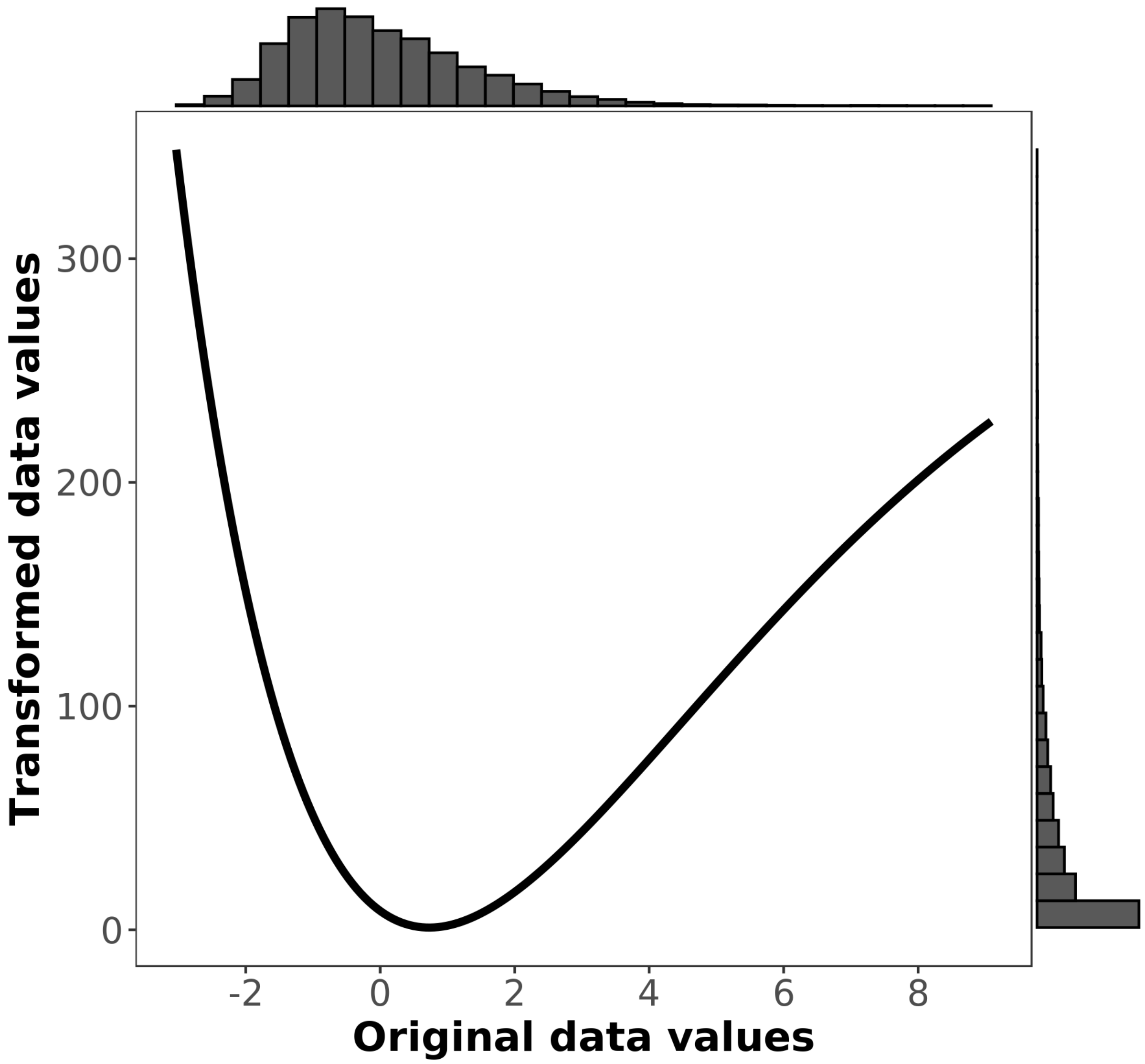


Figure S7. Resistance curve for PC axis 3, which explains 10.3% of the variation in the environment. High values are associated areas of high slope and ruggedness, while low values are associated with areas of low slope and ruggedness. Transformed data values indicate the estimated resistance for a given value along the PC axis.

### Yosemite

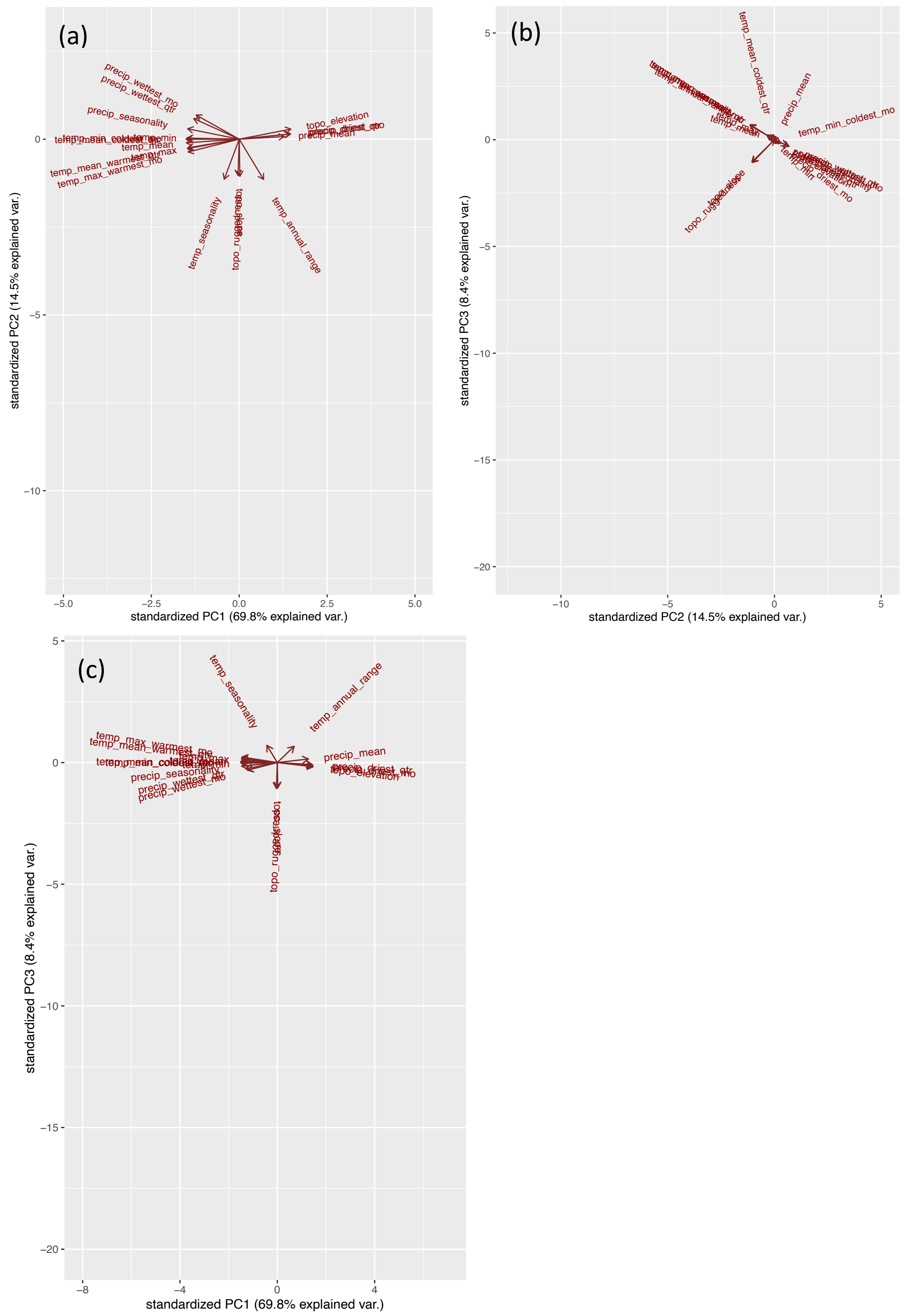


Figure S8. PCA plots of temperature, precipitation, and topography in the Yosemite region. Panel (a) shows component 1 vs component 2; panel (b) shows component 2 vs component 3; and panel (c) shows component 1 vs component 3.

Table S11. PCA loadings for component 1 in the Yosemite region, which explains 69.8% of the variation in the environment.

| Surface | PC1 |
|---|---|
| topo_elevation | 0.2711 |
| precip_driest_mo | 0.2697 |
| precip_driest_qtr | 0.2647 |
| precip_mean | 0.2360 |
| temp_annual_range | 0.1278 |
| topo_slope | 0.0032 |
| topo_ruggedness | -0.0064 |
| temp_seasonality | -0.0788 |
| precip_wettest_mo | -0.2261 |
| precip_wettest_qtr | -0.2388 |
| temp_max_warmest_mo | -0.2679 |
| temp_max | -0.2697 |
| precip_seasonality | -0.2705 |
| temp_min | -0.2718 |
| temp_mean_warmest_qtr | -0.2722 |
| temp_mean | -0.2772 |
| temp_mean_coldest_qtr | -0.2778 |
| temp_min_coldest_mo | -0.2785 |

Table S12. PCA loadings for component 2 in the Yosemite region, which explains 14.5% of the variation in the environment.

| Surface | PC2 |
|---|---|
| precip_wettest_mo | 0.2780 |
| precip_wettest_qtr | 0.2368 |
| precip_seasonality | 0.1197 |
| topo_elevation | 0.1161 |
| precip_driest_mo | 0.0608 |
| precip_driest_qtr | 0.0590 |
| precip_mean | 0.0304 |
| temp_min | 0.0126 |
| temp_min_coldest_mo | 0.0061 |
| temp_mean_coldest_qtr | -0.0010 |
| temp_mean | -0.0421 |
| temp_max | -0.0991 |
| temp_mean_warmest_qtr | -0.1091 |
| temp_max_warmest_mo | -0.1475 |
| topo_ruggedness | -0.4176 |
| topo_slope | -0.4285 |
| temp_seasonality | -0.4611 |
| temp_annual_range | -0.4634 |

Table S13. PCA loadings for component 3 in the Yosemite region, which explains 8.4% of the variation in the environment.

| Surface | PC3 |
|---|---|
| temp_seasonality | 0.3764 |
| temp_annual_range | 0.3558 |
| temp_max_warmest_mo | 0.1208 |
| temp_mean_warmest_qtr | 0.0900 |
| precip_mean | 0.0783 |
| temp_max | 0.0571 |
| temp_mean | 0.0194 |
| temp_mean_coldest_qtr | 0.0050 |
| temp_min_coldest_mo | 0.0021 |
| temp_min | -0.0164 |
| precip_driest_qtr | -0.0429 |
| precip_driest_mo | -0.0619 |
| precip_seasonality | -0.0798 |
| topo_elevation | -0.0917 |
| precip_wettest_qtr | -0.1373 |
| precip_wettest_mo | -0.1659 |
| topo_slope | -0.5548 |
| topo_ruggedness | -0.5695 |

Table S14. Jackknife results for landscape genetics analysis in the Yosemite region.

| surface | avg.AIC | avg.AICc | avg.weight | avg.rank | avg.R2m | avg.LL | n | Percent.top | k |
|---|---|---|---|---|---|---|---|---|---|
| Comp2 | -2833.416527 | -2831.511765 | 0.250194622 | 3.048 | 0.130001155 | 1420.708263 | 565 | 56.5 | 4 |
| Distance | -2830.917462 | -2830.395722 | 0.1080441 | 3.337 | 0.085104437 | 1419.458731 | 78 | 7.8 | 2 |
| Comp1 | -2832.137304 | -2830.232542 | 0.141916199 | 4.73 | 0.099140668 | 1420.068652 | 272 | 27.2 | 4 |
| roads | -2831.123244 | -2830.032335 | 0.089933521 | 4.805 | 0.086210213 | 1419.561622 | 0 | 0 | 3 |
| Comp3 | -2831.655026 | -2829.750264 | 0.07863238 | 5.399 | 0.086782389 | 1419.827513 | 3 | 0.3 | 4 |
| Comp2.roads | -2834.191155 | -2829.770102 | 0.088174785 | 5.413 | 0.128223503 | 1421.095577 | 7 | 0.7 | 6 |
| Comp2.Comp3 | -2835.673318 | -2829.451096 | 0.071197993 | 5.818 | 0.100161265 | 1421.836659 | 25 | 2.5 | 7 |
| Comp3.roads | -2833.625857 | -2829.204804 | 0.067537413 | 6.228 | 0.09332295 | 1420.812928 | 50 | 5 | 6 |
| Comp1.roads | -2832.305207 | -2827.884154 | 0.043986842 | 8.469 | 0.101182604 | 1420.152603 | 0 | 0 | 6 |
| Comp1.Comp3 | -2833.587388 | -2827.365166 | 0.025936732 | 9.585 | 0.090079057 | 1420.793694 | 0 | 0 | 7 |
| Comp1.Comp2 | -2833.30032 | -2827.078098 | 0.021301063 | 9.813 | 0.093985075 | 1420.65016 | 0 | 0 | 7 |
| Comp2.Comp3.roads | -2836.366284 | -2825.116284 | 0.008213551 | 11.546 | 0.112828142 | 1422.183142 | 0 | 0 | 9 |
| Comp1.Comp2.roads | -2834.019707 | -2822.769707 | 0.002578758 | 13.317 | 0.108162298 | 1421.009854 | 0 | 0 | 9 |
| Comp1.Comp3.roads | -2833.358855 | -2822.108855 | 0.001751873 | 13.609 | 0.088234271 | 1420.679427 | 0 | 0 | 9 |
| Comp1.Comp2.Comp3 | -2834.213223 | -2819.546557 | 0.000455502 | 14.959 | 0.10371053 | 1421.106612 | 0 | 0 | 10 |
| roads.vegetation | -2834.832573 | -2815.97543 | 9.21E-05 | 16.051 | 0.084586884 | 1421.416286 | 0 | 0 | 11 |
| Comp1.Comp2.Comp3.roads | -2836.695625 | -2812.695625 | 1.55E-05 | 17.329 | 0.109816193 | 1422.347813 | 0 | 0 | 12 |
| Comp2.vegetation | -2833.676378 | -2809.676378 | 5.90E-06 | 18.913 | 0.140488152 | 1420.838189 | 0 | 0 | 12 |
| Comp3.vegetation | -2832.983889 | -2808.983889 | 2.51E-06 | 19.277 | 0.079135365 | 1420.491945 | 0 | 0 | 12 |
| Comp1.vegetation | -2832.260708 | -2808.260708 | 3.72E-06 | 19.591 | 0.130117582 | 1420.130354 | 0 | 0 | 12 |
| vegetation | -2816.2478 | -2804.9978 | 2.50E-05 | 20.116 | 0.320428183 | 1412.1239 | 0 | 0 | 9 |
| Comp2.roads.vegetation | -2833.926733 | -2795.744915 | 6.42E-09 | 22.511 | 0.150247237 | 1420.963367 | 0 | 0 | 14 |
| Comp1.roads.vegetation | -2833.180181 | -2794.998363 | 2.87E-09 | 22.825 | 0.094530471 | 1420.590091 | 0 | 0 | 14 |
| Comp3.roads.vegetation | -2832.132161 | -2793.950343 | 1.33E-09 | 23.372 | 0.0808065 | 1420.066081 | 0 | 0 | 14 |
| Comp1.Comp2.vegetation | -2834.08512 | -2786.08512 | 3.19E-11 | 25.76 | 0.132515113 | 1421.04256 | 0 | 0 | 15 |
| Comp2.Comp3.vegetation | -2833.79181 | -2785.79181 | 4.41E-11 | 25.802 | 0.148048232 | 1420.895905 | 0 | 0 | 15 |

| | | | | | | | | | |
|---|---|---|---|---|---|---|---|---|---|
| Comp1.Comp3.vegetation | -2832.258176 | -2784.258176 | 1.63E-11 | 26.377 | 0.102584911 | 1420.129088 | 0 | 0 | 15 |
| Comp2.Comp3.roads.vegetation | -2833.854491 | -2757.354491 | 2.57E-17 | 28.844 | 0.149025174 | 1420.927245 | 0 | 0 | 17 |
| Comp1.Comp2.roads.vegetation | -2833.998157 | -2757.498157 | 1.99E-17 | 28.863 | 0.13239791 | 1420.999078 | 0 | 0 | 17 |
| Comp1.Comp3.roads.vegetation | -2832.844061 | -2756.344061 | 1.69E-17 | 29.293 | 0.104632859 | 1420.42203 | 0 | 0 | 17 |
| Comp1.Comp2.Comp3.vegetation | -2832.726806 | -2735.01252 | 2.66E-22 | 31 | 0.128278476 | 1420.363403 | 0 | 0 | 18 |
| Comp1.Comp2.Comp3.roads.vegetation | -2834.052446 | -2666.052446 | 2.63E-37 | 32 | 0.131867908 | 1421.026223 | 0 | 0 | 20 |

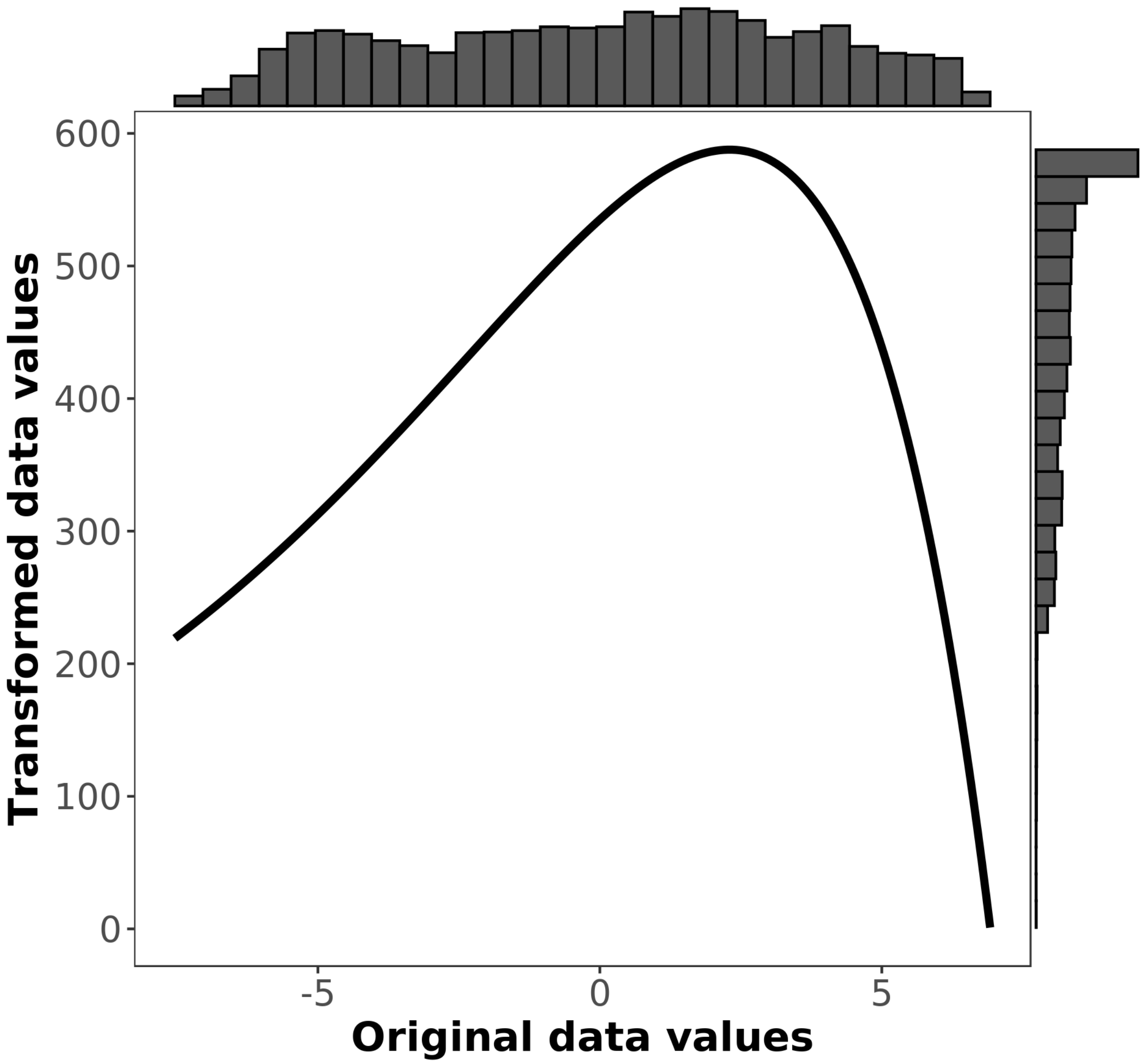


Figure S9. Resistance curve for PC axis 1, which explains 69.8% of the variation in the environment. High values are associated with higher elevation regions with more dry season precipitation and cooler temperatures, while low values are associated with lower elevation areas of higher temperature. Transformed data values indicate the estimated resistance for a given value along the PC axis.

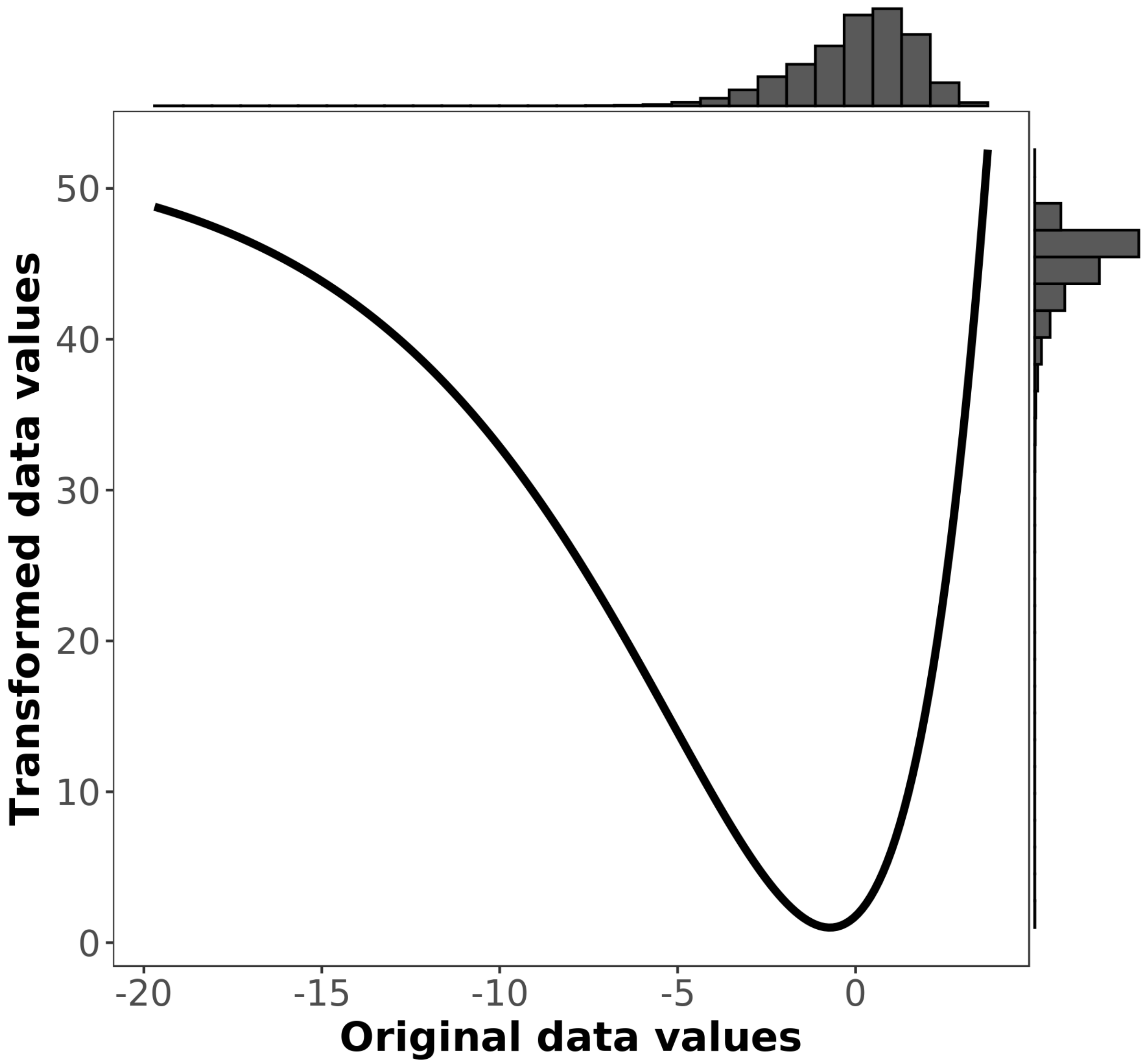


Figure S10. Resistance curve for PC axis 2, which explains 14.5% of the variation in the environment. Low values are associated with areas of higher topographic slope and ruggedness that also exhibit higher temperature seasonality and temperature annual range. Transformed data values indicate the estimated resistance for a given value along the PC axis.

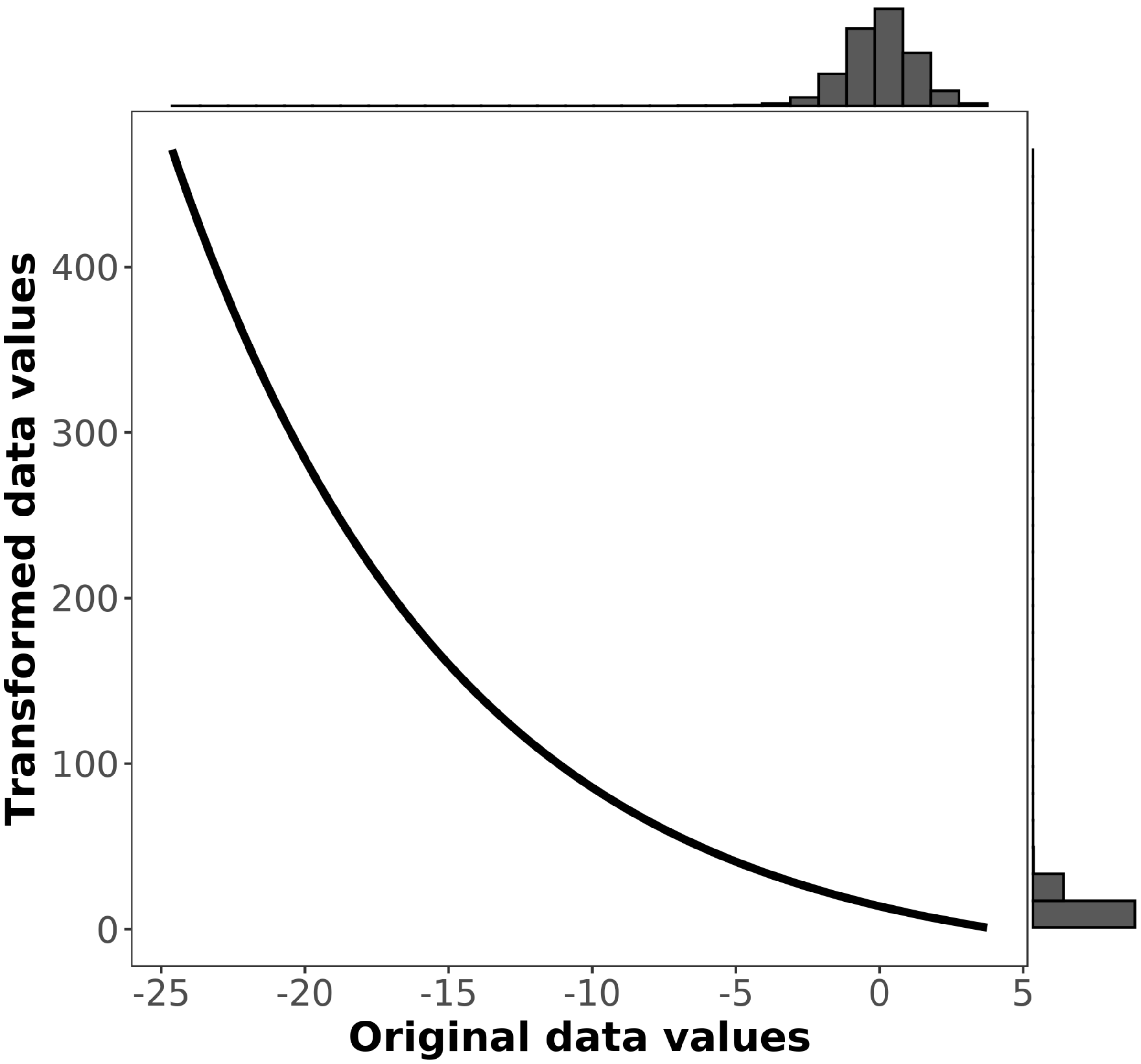


Figure S11. Resistance curve for PC axis 3, which explains 8.4% of the variation in the environment. High values are associated areas of high temperature seasonality and temperature annual range, while low values are associated with rugged areas of high slope. Transformed data values indicate the estimated resistance for a given value along the PC axis.

Table S15. Resistance value for roads in the Yosemite region.

| Feature | Resistance value |
| --- | --- |
| No road | 1 |
| Road | 499.88 |

### Tahoe

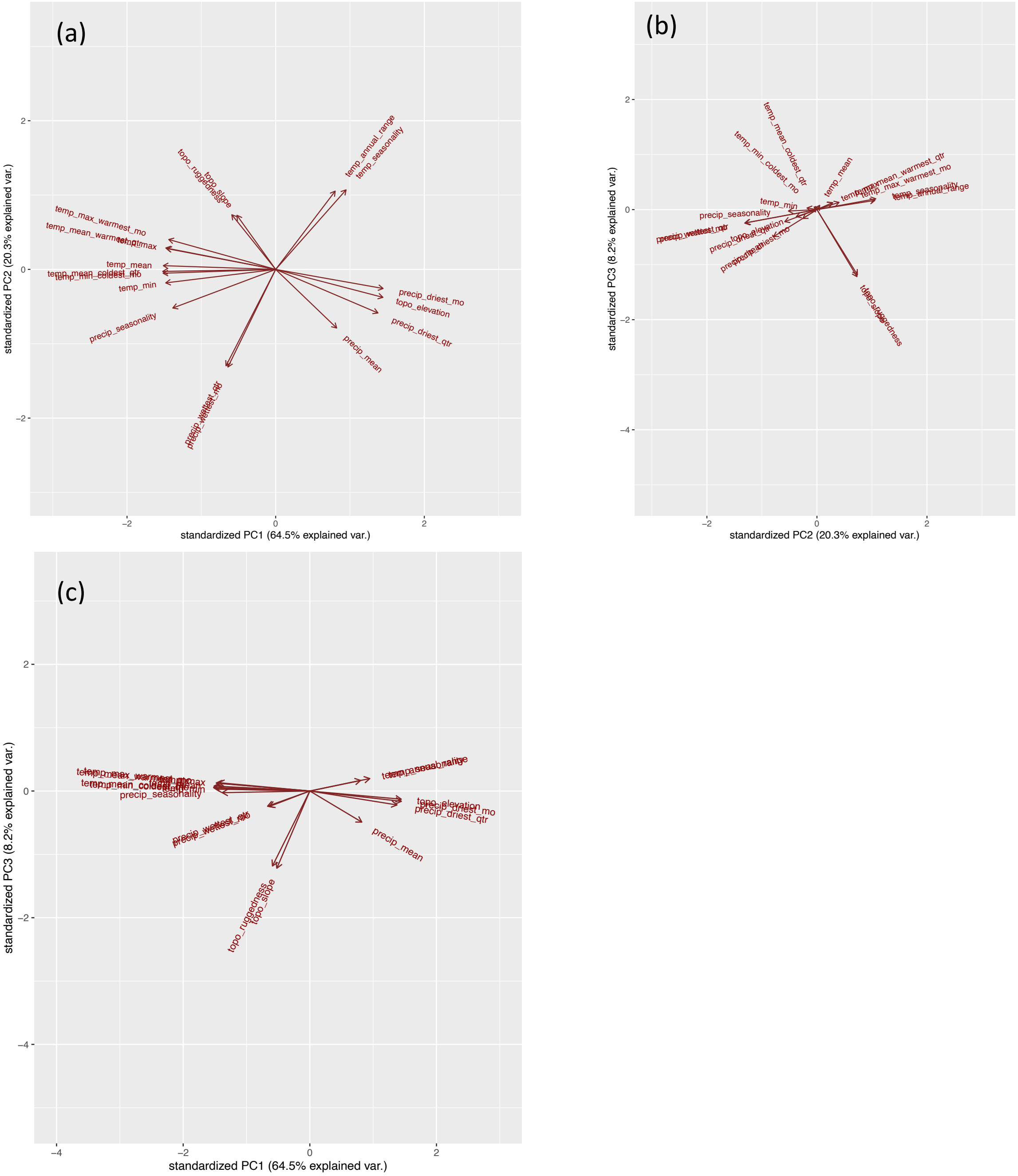


Figure S12. PCA plots of temperature, precipitation, and topography in the Tahoe region. Panel (a) shows component 1 vs component 2; panel (b) shows component 2 vs component 3; and panel (c) shows component 1 vs component 3.

Table S16. PCA loadings for component 1 in the Tahoe region, which explains 64.5% of the variation in the environment.

| Surface | PC1 |
|---|---|
| precip_driest_mo | 0.2774 |
| topo_elevation | 0.2771 |
| precip_driest_qtr | 0.2645 |
| temp_seasonality | 0.1817 |
| precip_mean | 0.1585 |
| temp_annual_range | 0.1547 |
| topo_slope | -0.0984 |
| topo_ruggedness | -0.1117 |
| precip_wettest_mo | -0.1222 |
| precip_wettest_qtr | -0.1272 |
| precip_seasonality | -0.2660 |
| temp_max_warmest_mo | -0.2763 |
| temp_max | -0.2811 |
| temp_mean_warmest_qtr | -0.2837 |
| temp_min | -0.2840 |
| temp_mean | -0.2901 |
| temp_min_coldest_mo | -0.2907 |
| temp_mean_coldest_qtr | -0.2917 |

Table S17. PCA loadings for component 2 in the Tahoe region, which explains 20.3% of the variation in the environment.

| Surface | PC2 |
|---|---|
| temp_seasonality | 0.3665 |
| temp_annual_range | 0.3595 |
| topo_ruggedness | 0.2521 |
| topo_slope | 0.2498 |
| temp_max_warmest_mo | 0.1367 |
| temp_mean_warmest_qtr | 0.0982 |
| temp_max | 0.0960 |
| temp_mean | 0.0178 |
| temp_mean_coldest_qtr | -0.0090 |
| temp_min_coldest_mo | -0.0172 |
| temp_min | -0.0610 |
| precip_driest_mo | -0.0865 |
| topo_elevation | -0.1286 |
| precip_seasonality | -0.1766 |
| precip_driest_qtr | -0.1996 |
| precip_mean | -0.2699 |
| precip_wettest_qtr | -0.4447 |
| precip_wettest_mo | -0.4504 |

Table S18. PCA loadings for component 3 in the Tahoe region, which explains 8.2% of the variation in the environment.

| Surface | PC3 |
|---|---|
| temp_seasonality | 0.1072 |
| temp_annual_range | 0.0886 |
| temp_max_warmest_mo | 0.0702 |
| temp_mean_warmest_qtr | 0.0674 |
| temp_max | 0.0650 |
| temp_mean | 0.0414 |
| temp_mean_coldest_qtr | 0.0305 |
| temp_min_coldest_mo | 0.0266 |
| temp_min | 0.0157 |
| precip_seasonality | -0.0144 |
| topo_elevation | -0.0685 |
| precip_driest_mo | -0.0867 |
| precip_driest_qtr | -0.1171 |
| precip_wettest_qtr | -0.1310 |
| precip_wettest_mo | -0.1335 |
| precip_mean | -0.2622 |
| topo_ruggedness | -0.6362 |
| topo_slope | -0.6548 |

Table S19. Jackknife results for landscape genetics analysis in the Tahoe region.

| surface | avg.AIC | avg.AICc | avg.weight | avg.rank | avg.R2m | avg.LL | n | Percent.top | k |
|---|---|---|---|---|---|---|---|---|---|
| Comp2 | -4177.289176 | -4175.622509 | 0.328596867 | 1.671 | 0.016578358 | 2092.644588 | 688 | 68.8 | 4 |
| Comp3 | -4175.574854 | -4173.908187 | 0.165300216 | 3.087 | 0.01436222 | 2091.787427 | 229 | 22.9 | 4 |
| Comp2.roads | -4177.246655 | -4173.428474 | 0.118404116 | 4.003 | 0.019938987 | 2092.623328 | 1 | 0.1 | 6 |
| Comp1 | -4173.44886 | -4171.782194 | 0.070346139 | 5.662 | 0.017343004 | 2090.72443 | 48 | 4.8 | 4 |
| Comp1.Comp2 | -4177.537306 | -4172.203973 | 0.066342909 | 5.847 | 0.015142931 | 2092.768653 | 7 | 0.7 | 7 |
| Comp3.roads | -4175.491973 | -4171.673791 | 0.053283425 | 6.204 | 0.014421811 | 2091.745986 | 0 | 0 | 6 |
| Comp2.Comp3 | -4177.314619 | -4171.981286 | 0.05176589 | 6.298 | 0.016601119 | 2092.657309 | 0 | 0 | 7 |
| Comp1.Comp3 | -4175.503257 | -4170.169924 | 0.025157592 | 8.709 | 0.014408047 | 2091.751629 | 0 | 0 | 7 |
| Comp1.roads | -4173.823565 | -4170.005383 | 0.030170771 | 8.816 | 0.018743599 | 2090.911782 | 0 | 0 | 6 |
| roads | -4170.46631 | -4169.50631 | 0.035853762 | 8.966 | 0.016076771 | 2089.233155 | 2 | 0.2 | 3 |
| Distance | -4169.605132 | -4169.143594 | 0.039743337 | 9.372 | 0.015697638 | 2088.802566 | 25 | 2.5 | 2 |
| Comp1.Comp2.roads | -4177.061778 | -4167.588094 | 0.005389273 | 11.555 | 0.016513985 | 2092.530889 | 0 | 0 | 9 |
| Comp2.Comp3.roads | -4176.776908 | -4167.303224 | 0.004757031 | 12.121 | 0.015940049 | 2092.388454 | 0 | 0 | 9 |
| Comp1.Comp3.roads | -4175.387415 | -4165.913731 | 0.002951461 | 13.479 | 0.014507324 | 2091.693708 | 0 | 0 | 9 |
| Comp1.Comp2.Comp3 | -4177.304693 | -4165.08247 | 0.001578748 | 14.364 | 0.017938153 | 2092.652346 | 0 | 0 | 10 |
| vegetation | -4172.754967 | -4160.532745 | 0.00031153 | 16.064 | 0.020735993 | 2090.377484 | 0 | 0 | 10 |
| Comp1.Comp2.Comp3.roads | -4176.559289 | -4157.059289 | 2.89E-05 | 16.983 | 0.015624427 | 2092.279644 | 0 | 0 | 12 |
| roads.vegetation | -4173.173666 | -4153.673666 | 1.15E-05 | 18.384 | 0.021857833 | 2090.586833 | 0 | 0 | 12 |
| Comp2.vegetation | -4176.798668 | -4152.532001 | 3.60E-06 | 19.066 | 0.01591399 | 2092.399334 | 0 | 0 | 13 |
| Comp3.vegetation | -4175.265636 | -4150.99897 | 1.66E-06 | 19.855 | 0.014612659 | 2091.632818 | 0 | 0 | 13 |
| Comp1.vegetation | -4173.579358 | -4149.312691 | 1.26E-06 | 20.525 | 0.020694344 | 2090.789679 | 0 | 0 | 13 |
| Comp2.roads.vegetation | -4176.626842 | -4139.703765 | 4.99E-09 | 22.378 | 0.01556188 | 2092.313421 | 0 | 0 | 15 |
| Comp3.roads.vegetation | -4175.299507 | -4138.37643 | 3.15E-09 | 23.096 | 0.014394237 | 2091.649753 | 0 | 0 | 15 |
| Comp1.roads.vegetation | -4173.50195 | -4136.578873 | 1.78E-09 | 23.603 | 0.01908789 | 2090.750975 | 0 | 0 | 15 |
| Comp1.Comp2.vegetation | -4176.737318 | -4131.403985 | 7.99E-11 | 25.538 | 0.015890759 | 2092.368659 | 0 | 0 | 16 |
| Comp2.Comp3.vegetation | -4176.647039 | -4131.313706 | 8.17E-11 | 25.578 | 0.018900036 | 2092.323519 | 0 | 0 | 16 |

| | | | | | | | | | |
|---|---|---|---|---|---|---|---|---|---|
| Comp1.Comp3.vegetation | -4173.674141 | -4128.340808 | 2.30E-11 | 26.776 | 0.017005493 | 2090.837071 | 0 | 0 | 16 |
| Comp2.Comp3.roads.vegetation | -4176.76927 | -4108.36927 | 8.47E-16 | 28.284 | 0.015422741 | 2092.384635 | 0 | 0 | 18 |
| Comp1.Comp2.roads.vegetation | -4176.284498 | -4107.884498 | 6.29E-16 | 29.106 | 0.015448115 | 2092.142249 | 0 | 0 | 18 |
| Comp1.Comp3.roads.vegetation | -4174.83078 | -4106.43078 | 3.14E-16 | 29.61 | 0.015660235 | 2091.41539 | 0 | 0 | 18 |
| Comp1.Comp2.Comp3.vegetation | -4175.816907 | -4091.372463 | 1.55E-19 | 31 | 0.015470791 | 2091.908454 | 0 | 0 | 19 |
| Comp1.Comp2.Comp3.roads.vegetation | -4175.703306 | -4043.703306 | 7.27E-30 | 32 | 0.016595832 | 2091.851653 | 0 | 0 | 21 |

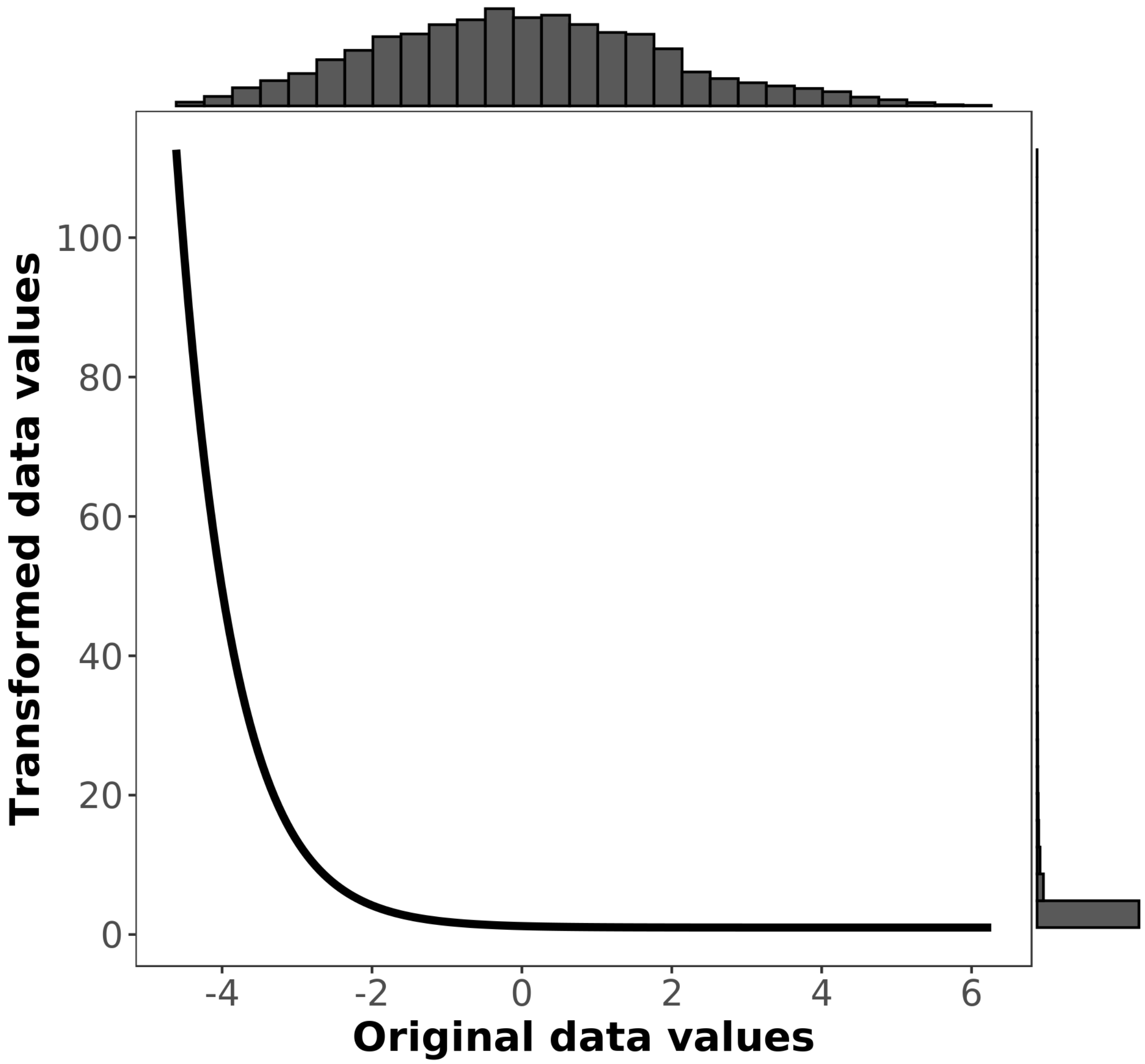


Figure S13. Resistance curve for PC axis 2, which explains 20.3% of the variation in the environment. High values are associated with areas of high temperature seasonality, high temperature annual range and rugged areas of high slope, while low values are associated with areas that receive more precipitation—especially wet season precipitation. Transformed data values indicate the estimated resistance for a given value along the PC axis.

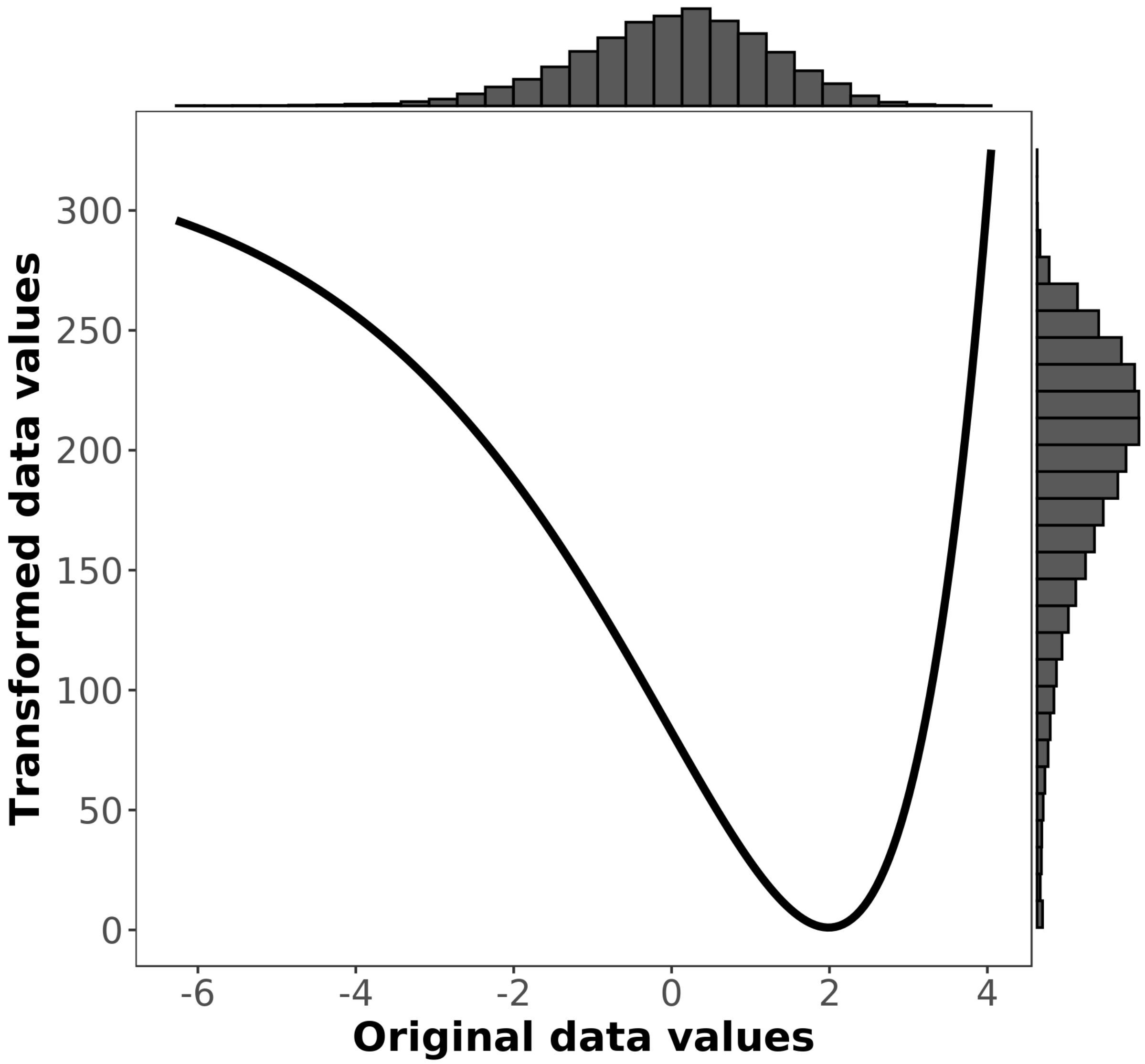


Figure S14. Resistance curve for PC axis 3, which explains 8.2% of the variation in the environment. High values are those with little topographic relief and low slope, while low values are associated with rugged areas of high slope. Transformed data values indicate the estimated resistance for a given value along the PC axis.

**Plumas**

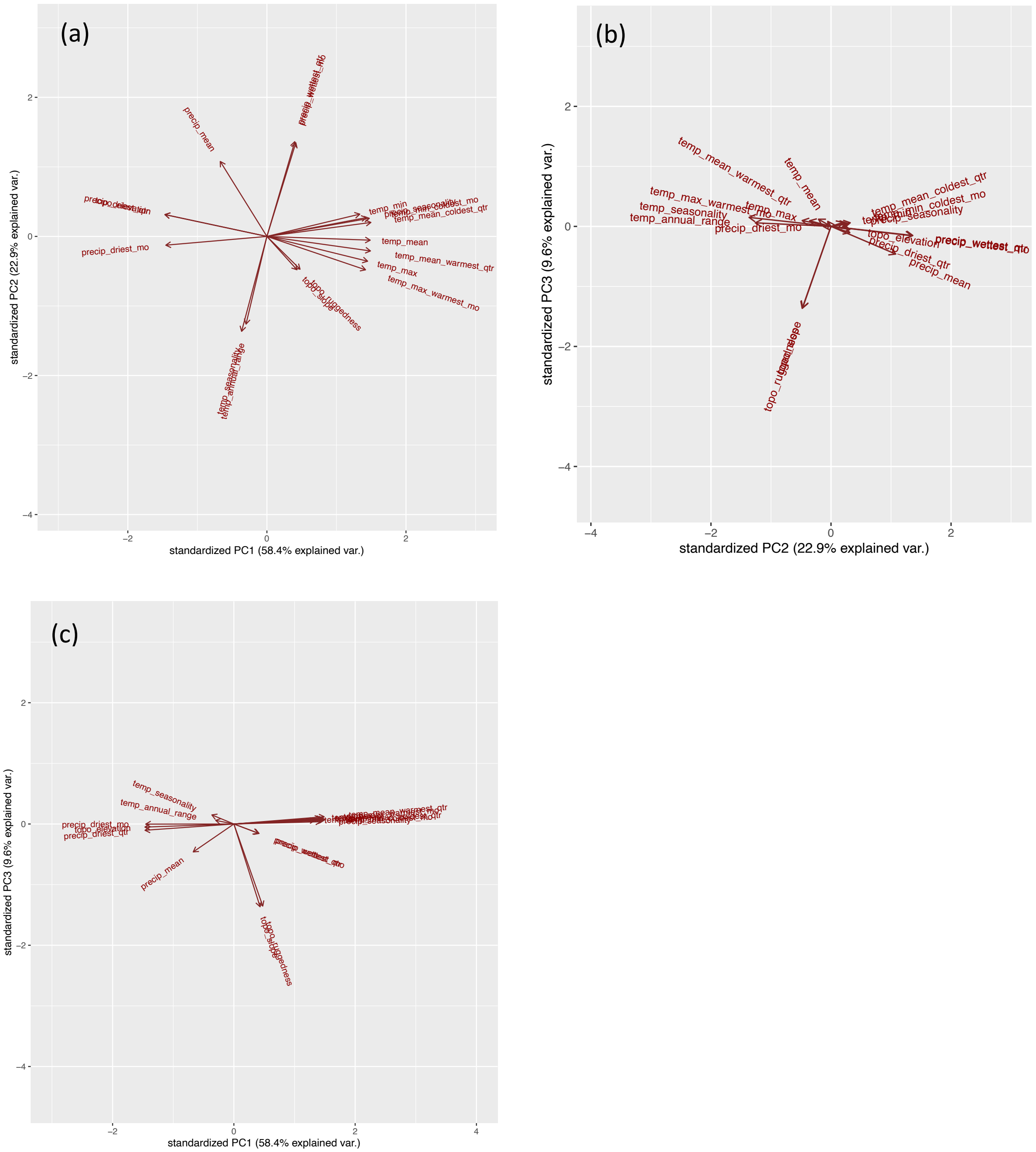


Figure S15. PCA plots of temperature, precipitation, and topography in the Plumas region. Panel (a) shows component 1 vs component 2; panel (b) shows component 2 vs component 3; and panel (c) shows component 1 vs component 3.

Table S20. PCA loadings for component 1 in the Tahoe region, which explains 58.4% of the variation in the environment.

| Surface | PC1 |
|---|---|
| temp_mean_coldest_qtr | 0.3015 |
| temp_mean_warmest_qtr | 0.3002 |
| temp_mean | 0.2995 |
| temp_min_coldest_mo | 0.2969 |
| temp_max | 0.2920 |
| precip_seasonality | 0.2885 |
| temp_max_warmest_mo | 0.2859 |
| temp_min | 0.2693 |
| topo_ruggedness | 0.0961 |
| topo_slope | 0.0872 |
| precip_wettest_mo | 0.0838 |
| precip_wettest_qtr | 0.0807 |
| temp_annual_range | -0.0606 |
| temp_seasonality | -0.0734 |
| precip_mean | -0.1352 |
| precip_driest_mo | -0.2931 |
| precip_driest_qtr | -0.2955 |
| topo_elevation | -0.2958 |

Table S21. PCA loadings for component 2 in the Tahoe region, which explains 22.9% of the variation in the environment.

| Surface | PC2 |
|---|---|
| precip_wettest_qtr | 0.4389 |
| precip_wettest_mo | 0.4339 |
| precip_mean | 0.3470 |
| temp_min | 0.1032 |
| topo_elevation | 0.1012 |
| precip_driest_qtr | 0.1010 |
| precip_seasonality | 0.0873 |
| temp_min_coldest_mo | 0.0838 |
| temp_mean_coldest_qtr | 0.0648 |
| temp_mean | -0.0174 |
| precip_driest_mo | -0.0405 |
| temp_mean_warmest_qtr | -0.0673 |
| temp_max | -0.1148 |
| topo_ruggedness | -0.1520 |
| topo_slope | -0.1545 |
| temp_max_warmest_mo | -0.1547 |
| temp_annual_range | -0.4061 |
| temp_seasonality | -0.4396 |

Table S22. PCA loadings for component 3 in the Tahoe region, which explains 9.6% of the variation in the environment.

| Surface | PC3 |
|---|---|
| temp_seasonality | 0.0747 |
| temp_mean_warmest_qtr | 0.0596 |
| temp_max_warmest_mo | 0.0484 |
| temp_max | 0.0468 |
| temp_mean | 0.0417 |
| temp_mean_coldest_qtr | 0.0336 |
| temp_min | 0.0301 |
| temp_annual_range | 0.0296 |
| temp_min_coldest_mo | 0.0280 |
| precip_seasonality | 0.0185 |
| precip_driest_mo | -0.0009 |
| topo_elevation | -0.0253 |
| precip_driest_qtr | -0.0515 |
| precip_wettest_qtr | -0.0752 |
| precip_wettest_mo | -0.0777 |
| precip_mean | -0.2289 |
| topo_ruggedness | -0.6729 |
| topo_slope | -0.6785 |

Table S23. Jackknife results for landscape genetics analysis in in the Plumas region.

| surface | avg.AIC | avg.AICc | avg.weight | avg.rank | avg.R2m | avg.LL | n | Percent.top | k |
|---|---|---|---|---|---|---|---|---|---|
| Comp1.Comp3.roads | -344.2043328 | -524.2043328 | 0.360465563 | 1.768 | 0.168007874 | 176.1021664 | 472 | 47.2 | 9 |
| Comp1.Comp2.roads | -344.472292 | -524.472292 | 0.37840758 | 1.874 | 0.138923757 | 176.236146 | 282 | 28.2 | 9 |
| Comp2.Comp3.roads | -343.1914644 | -523.1914644 | 0.261126856 | 2.358 | 0.229309263 | 175.5957322 | 246 | 24.6 | 9 |
| Comp1.Comp2.Comp3 | -344.208083 | -454.208083 | 2.25E-16 | 4 | 0.166666407 | 176.1040415 | 0 | 0 | 10 |
| vegetation | -341.0549782 | -429.0549782 | 1.11E-21 | 5 | 0.100270128 | 174.5274891 | 0 | 0 | 11 |
| Comp1.Comp2.Comp3.roads | -343.3400183 | -421.3400183 | 1.56E-23 | 6.009 | 0.132801731 | 175.6700092 | 0 | 0 | 12 |
| Comp1.Comp2.Comp3.roads.vegetation | -344.5545671 | -416.8402814 | 1.60E-24 | 7.082 | 0.148323921 | 176.2772835 | 0 | 0 | 22 |
| Comp1.vegetation | -344.5369202 | -414.5369202 | 5.53E-25 | 9.373 | 0.160190631 | 176.2684601 | 0 | 0 | 14 |
| Comp1.Comp2.Comp3.vegetation | -344.2688547 | -414.2688547 | 4.98E-25 | 10.453 | 0.170980176 | 176.1344273 | 0 | 0 | 20 |
| roads.vegetation | -341.0470054 | -413.8470054 | 5.55E-25 | 10.915 | 0.099828119 | 174.5235027 | 0 | 0 | 13 |
| Comp3.vegetation | -343.4881126 | -413.4881126 | 4.22E-25 | 11.861 | 0.265103363 | 175.7440563 | 0 | 0 | 14 |
| Comp1.Comp2.roads.vegetation | -344.5266926 | -413.6176017 | 3.29E-25 | 12.131 | 0.16405669 | 176.2633463 | 0 | 0 | 19 |
| Comp1.Comp3.roads.vegetation | -344.3903773 | -413.4812864 | 3.29E-25 | 12.533 | 0.172322808 | 176.1951887 | 0 | 0 | 19 |
| Comp2.Comp3.roads.vegetation | -343.1066995 | -412.1976086 | 2.11E-25 | 14.863 | 0.23168574 | 175.5533498 | 0 | 0 | 19 |
| Comp1.Comp2.vegetation | -344.6379292 | -412.6379292 | 1.97E-25 | 15.682 | 0.161603541 | 176.3189646 | 0 | 0 | 17 |
| Comp1.roads.vegetation | -344.5248683 | -412.5248683 | 2.00E-25 | 15.77 | 0.17343177 | 176.2624341 | 0 | 0 | 16 |
| Comp1.Comp3.vegetation | -344.4135667 | -412.4135667 | 1.95E-25 | 16.066 | 0.178938279 | 176.2067833 | 0 | 0 | 17 |
| Comp2.vegetation | -341.6338695 | -411.6338695 | 1.41E-25 | 16.946 | 0.092232513 | 174.8169348 | 0 | 0 | 14 |
| Comp3.roads.vegetation | -343.282128 | -411.282128 | 1.40E-25 | 17.51 | 0.256144412 | 175.641064 | 0 | 0 | 16 |
| Comp2.Comp3.vegetation | -343.1742331 | -411.1742331 | 1.26E-25 | 18.096 | 0.231557291 | 175.5871165 | 0 | 0 | 17 |
| Comp2.roads.vegetation | -341.5130644 | -409.5130644 | 4.95E-26 | 20.71 | 0.093491108 | 174.7565322 | 0 | 0 | 16 |
| Distance | -340.5061823 | -338.5061823 | 2.46E-41 | 22.099 | 0.059442542 | 174.2530911 | 0 | 0 | 2 |
| roads | -340.5078975 | -335.7078975 | 6.08E-42 | 23.339 | 0.059430298 | 174.2539488 | 0 | 0 | 3 |
| Comp1 | -344.2622436 | -334.2622436 | 2.12E-42 | 24.129 | 0.172777522 | 176.1311218 | 0 | 0 | 4 |
| Comp3 | -343.3902131 | -333.3902131 | 1.71E-42 | 24.715 | 0.259031487 | 175.6951065 | 0 | 0 | 4 |
| Comp2 | -341.7566607 | -331.7566607 | 7.58E-43 | 25.718 | 0.22769781 | 174.8783303 | 0 | 0 | 4 |

| | | | | | | | | | |
|---|---|---|---|---|---|---|---|---|---|
| Comp1.roads | -344.2509917 | -302.2509917 | 2.30E-49 | 27.448 | 0.169504774 | 176.1254958 | 0 | 0 | 6 |
| Comp3.roads | -343.2236567 | -301.2236567 | 1.77E-49 | 27.785 | 0.246642655 | 175.6118283 | 0 | 0 | 6 |
| Comp2.roads | -341.5766927 | -299.5766927 | 7.47E-50 | 28.767 | 0.223805348 | 174.7883463 | 0 | 0 | 6 |
| Comp1.Comp3 | -344.2499997 | -232.2499997 | 1.45E-64 | 30.785 | 0.16945549 | 176.1249999 | 0 | 0 | 7 |
| Comp1.Comp2 | -344.420439 | -232.420439 | 1.44E-64 | 30.96 | 0.157050208 | 176.2102195 | 0 | 0 | 7 |
| Comp2.Comp3 | -343.3860113 | -231.3860113 | 1.13E-64 | 31.255 | 0.258333073 | 175.6930057 | 0 | 0 | 7 |

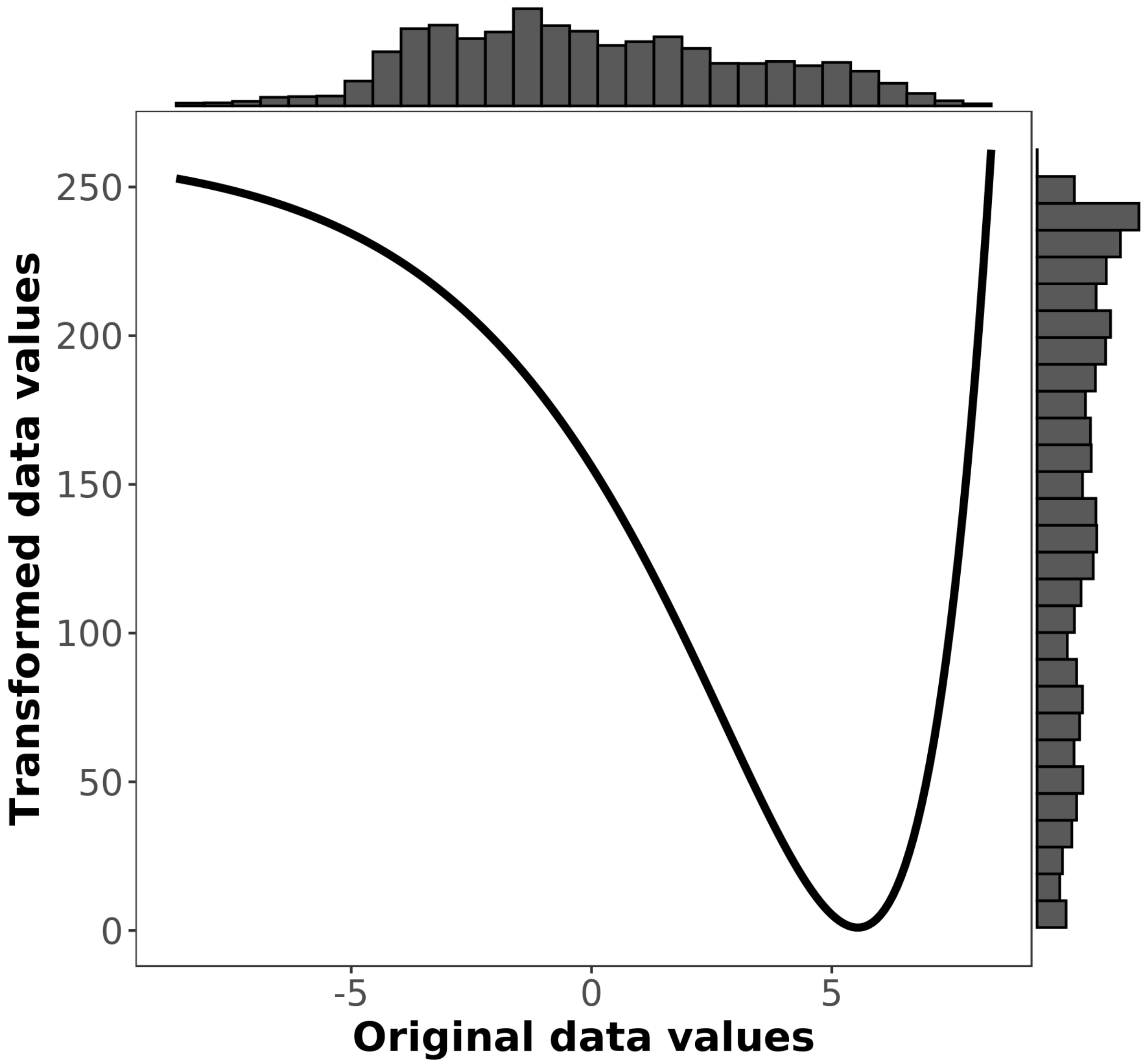


Figure S16. Resistance curve for PC axis 1, which explains 58.4% of the variation in the environment. High values are associated with warmer low-elevation areas, while low values are associated with colder high-elevation areas. Transformed data values indicate the estimated resistance for a given value along the PC axis.

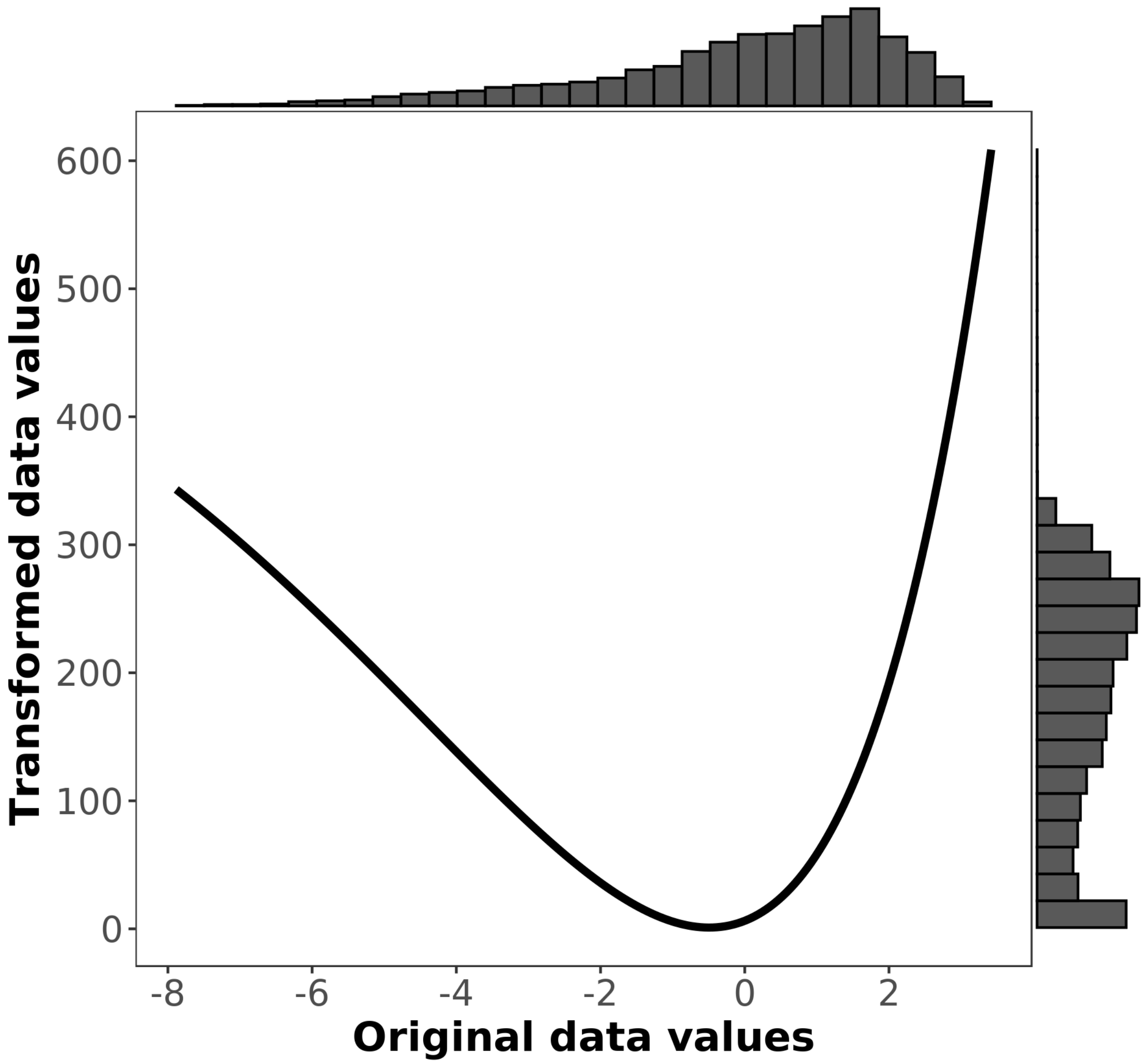


Figure S17. Resistance curve for PC axis 2, which explains 22.9% of the variation in the environment. High values are associated with wetter areas (especially wet-season precipitation) while low values are associated with more temperature seasonality. Transformed data values indicate the estimated resistance for a given value along the PC axis.

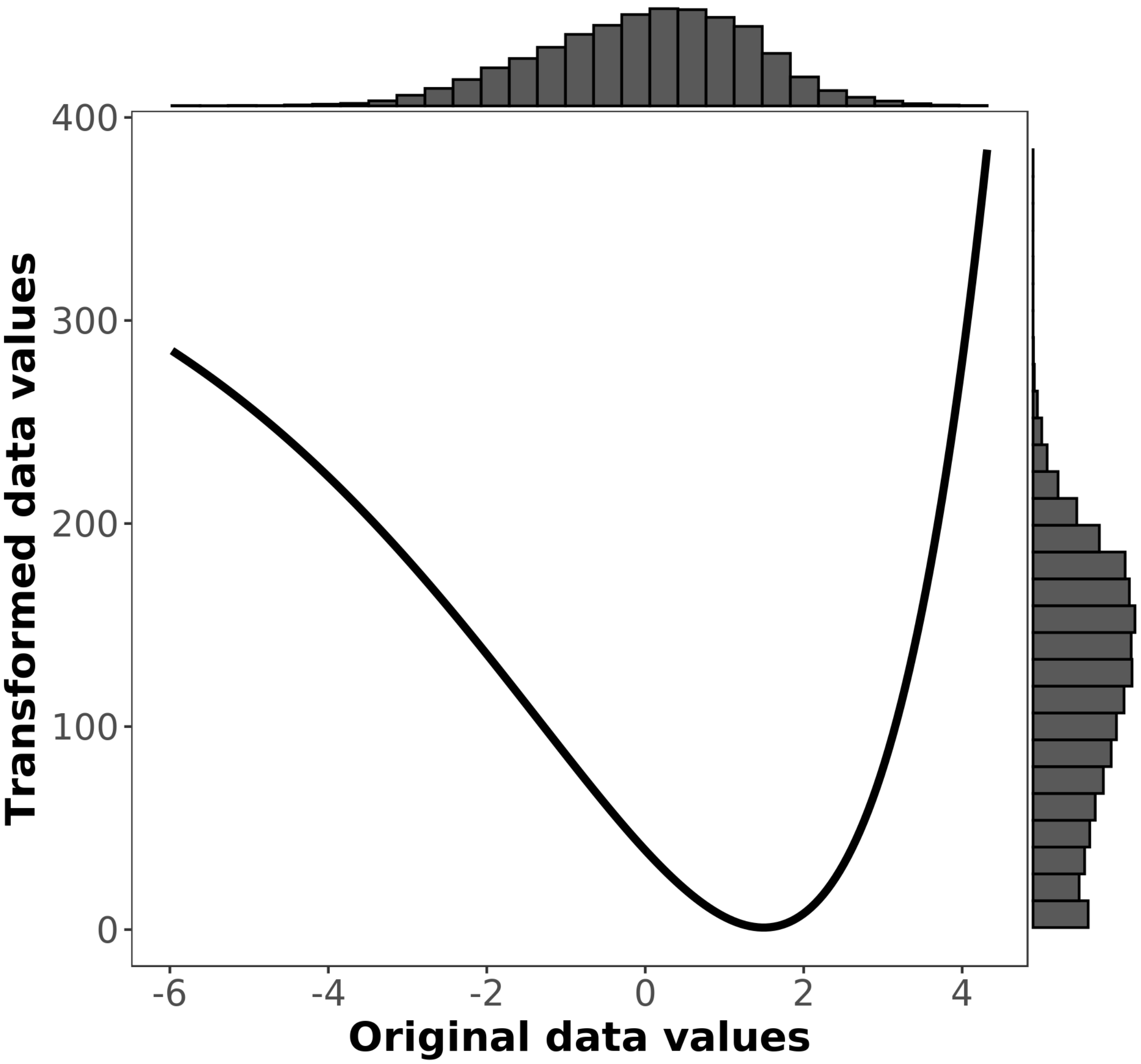


Figure S18. Resistance curve for PC axis 3, which explains 9.6% of the variation in the environment. High values have little topographic relief and low slope, while low values are associated with rugged areas of high slope. Transformed data values indicate the estimated resistance for a given value along the PC axis.

Table S24. Resistance value for roads in the Plumas region.

| Feature | Resistance value |
|---|---|
| No road | 2.27 |
| Road | 1 |
